# A generalizable framework to comprehensively predict epigenome, chromatin organization, and transcriptome

**DOI:** 10.1101/2022.05.23.493129

**Authors:** Zhenhao Zhang, Fan Feng, Yiyang Qiu, Jie Liu

## Abstract

Many deep learning approaches have been proposed to predict epigenetic profiles, chromatin organization, and transcription activity. While these approaches achieve satisfactory performance in predicting one modality from another, the learned representations are not generalizable across predictive tasks or across cell types. In this paper, we propose a deep learning approach named EPCOT which employs a pre-training and fine-tuning framework, and comprehensively predicts epigenome, chromatin organization, transcriptome, and enhancer activity in one framework. EPCOT is the first framework proposed to predict all of these genomic modalities and performs well in individual modality prediction, which is also generalizable to new cell and tissue types. EPCOT also maps from DNA sequence and chromatin accessibility profiles to generic representations which are generalizable across different modalities. Interpreting EPCOT model also provides biological insights including mapping between different genomic modalities, identifying TF sequence binding patterns, and analyzing cell-type specific TF impacts on enhancer activity.

## Main

Recent computational models have shown great promise in capturing the connection among human genome, epigenome, 3D chromatin organization, and transcriptome at a genome-wide scale. Some of them predict epigenomic features from DNA sequence [1, 2, 3, 4]. Some of them predict gene expression [3, 5, 6, 7], high-resolution 3D chromatin contact maps [8, 9, 10, 11], and enhancer activity [12, 13, 14] from DNA sequence or from epigenomic feature profiles (Extended Data Fig.1). However, these models typically predict one modality from another modality separately. To the best of our knowledge, there is no computational framework that is trained in an end-to-end fashion to comprehensively predict epigenome, 3D chromatin organization, and transcriptome. Another issue with many of these computational models is the obscure of the cell type and tissue type specificity. Some of these models usually make predictions from the DNA sequence and are trained with multiple cell types altogether. As a result, the trained model itself is not specific to any cell and tissue type. While some models [2, 7, 10] can predict for new cell types, it is unclear whether the models, representations, and insights learned are generalizable from one predictive task to another.

In this work, we propose EPCOT (comprehensively predicting <u>EP</u>igenome, <u>C</u>hromatin <u>O</u>rganization and <u>T</u>ranscription), a framework to comprehensively predict epigenomic features, gene expression, chromatin contact maps, and enhancer activity from DNA sequence and cell-type specific chromatin accessibility profiles (Fig.1a). The model leverages a popular pre-training and fine-tuning framework. The pre-training model has an encoder-decoder structure and performs epigenomic feature prediction (EFP). The encoder learns sequence representations from the inputs, whereas the decoder captures dependence among epigenomic features and selects sequence representations of interest to predict the epigenomic features. In the fine-tuning stage, the sequence representations yielded from pre-training model’s encoder are transferred to downstream tasks including gene expression prediction (GEP), chromatin organization prediction (COP), and enhancer activity prediction (EAP). Compared with multiple sequence-based or epigenomic feature-based predictive models, EPCOT achieves superior predictive performance or comparable performance by only using chromatin accessibility and reference sequence in all the tasks. Additionally, different from models that are pre-trained and fine-tuned on DNA sequence only [9, 15], EPCOT learns cell-type specific representations from DNA sequence with chromatin accessibility signals, which is generalizable to new, unseen human cell and tissue types by only requiring cell-type specific chromatin accessibility data as input. Here, chromatin accessibility is leveraged to provide cell-type specific information due to its connection with TF binding, gene regulation, and chromatin contacts [16, 17, 18]. Furthermore, EPCOT trained on human cell types can be generalized to mouse cell and tissue types. EPCOT also learns generic sequence representations which are generalizable among different predictive tasks. In addition, EPCOT is interpretable and reveals biological insights. First, EPCOT characterizes transcription factor (TF) co-binding patterns as reflected in learned TF embeddings, and also identifies TF sequence binding patterns that associate with their binding motifs, co-binding, or some unknown sequence patterns, which will help researchers further understand TF binding mechanisms. Second, EPCOT captures cell-type specific relationships between epigenomic features and transcriptional activity. Especially, EPCOT identifies differential contributions of epigenomic features in GEP and EAP tasks.

**Fig.1:**
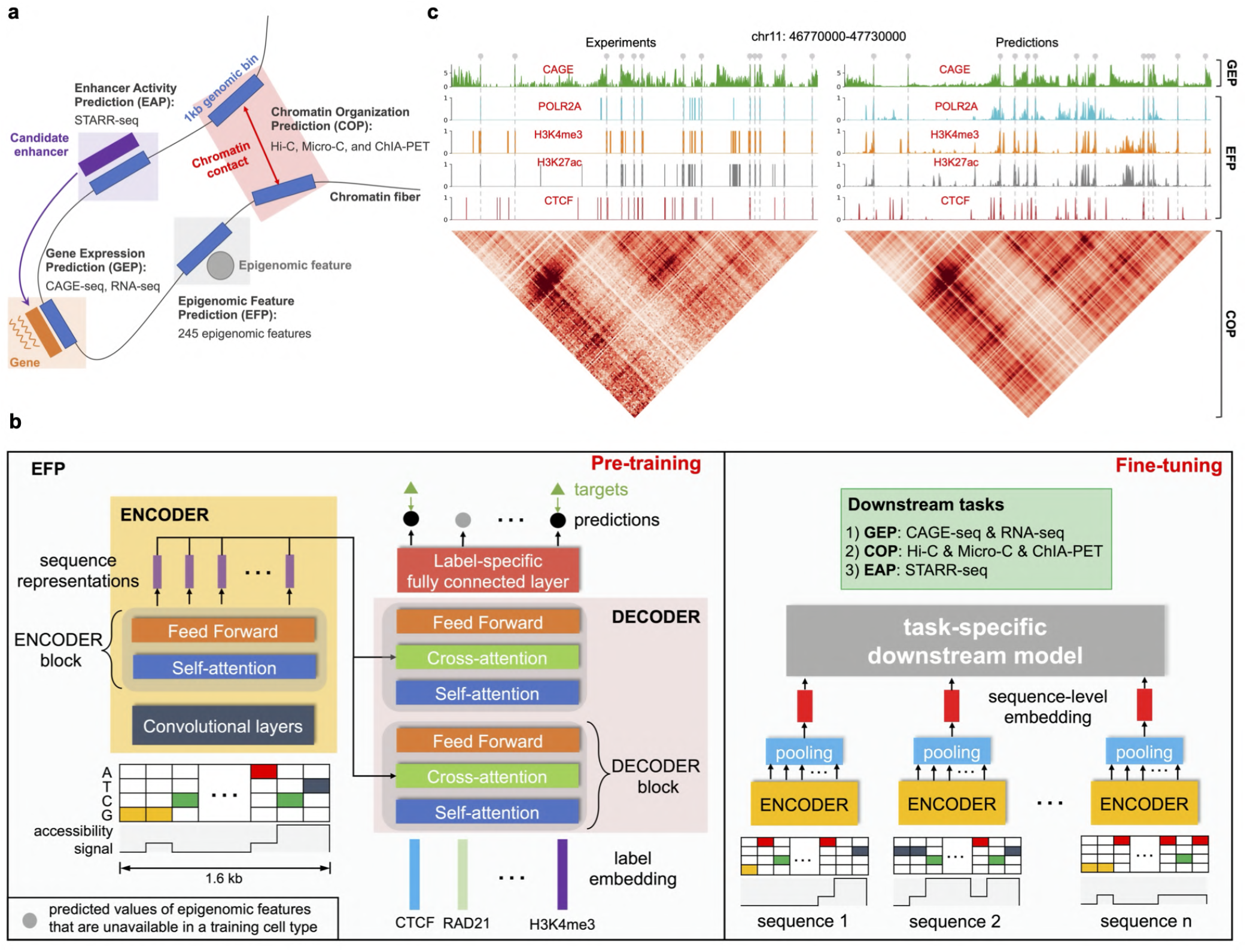
A pre-training and fine-tuning framework to comprehensively predict multiple genomic modalities. **a**, EPCOT predicts four modalities including EFP, GEP, COP, and EAP. **b**, Two-stage training of our EPCOT model including pre-training (left) and fine-tuning (right). The pre-training EFP model takes the inputs of 1.6kb genomic sequences including the 300bp flanking regions upstream and downstream of a 1kb region and its chromatin accessibility signal. In the fine-tuning stage, a task-specific downstream model is built on the pre-training model’s encoder with the inputs of multiple genomic sequences, and the downstream model is trained to complete GEP, COP, and EAP tasks. **c**, EPCOT successfully predicts epigenomic features, CAGE-seq, and Hi-C contact maps. In a 960kb example region in GM12878, CAGE-seq, POLR2A, H3K4me3, and H3K27ac profiles as well as Hi-C contact maps from EPCOT and the targets are compared.

Our work contributes novel and rigorous computational methods for scientific discoveries from large-scale genomic, epigenomic and transcriptomic datasets, and elucidates the new understanding of human genome functions from a comprehensive perspective. To our best knowledge, EPCOT is the first computational framework that comprehensively predicts extensive genomic modalities in a cell-type specific manner. As one of the final products in our framework, the generic embeddings of different genomic entities can be used in other large-scale computational and predictive models as the generic embeddings include information relevant to the epigenome, 3D chromatin organization, and transcriptional activity. The mapping from DNA sequence and chromatin accessibility profiles to numerical vectors allows the computers to use a universal language to predict these genomic modalities.

## Results

### A pre-training and fine-tuning framework to predict detailed epigenome, high-resolution chromatin organization, transcription, and enhancer activity from cell-type specific chromatin accessibility profiles

Our deep learning model EPCOT uses a two-stage pre-training and fine-tuning framework. In the first pretraining stage (Fig.1b left), a supervised pre-training model is designed to predict 245 epigenomic features including 236 TFs and 9 histone modifications, from the inputs of genomic sequence and its cell-type specific chromatin accessibility profile. The whole DNA sequence is segmented into 1kb central genomic sequences, and 300bp flanking regions both upstream and downstream of each 1kb sequence are appended, which results in 1.6kb short sequences. EPCOT pre-trains on each short genomic sequence. The pre-training model uses a unique encoder-decoder framework [19, 20]. The encoder learns sequence representations of the inputs, which are used in downstream tasks. The decoder learns the dependence among the epigenomic features and combines sequence representations to predict epigenomic features.

The fine-tuning stage (Fig.1b right) includes a number of downstream tasks related to epigenomic features including high-resolution chromatin organization (measured by Hi-C, Micro-C, and ChIA-PET), transcription (measured by CAGE-seq and RNA-seq) and enhancer activity (measured by STARR-seq). The downstream tasks are typically completed on longer genomic sequences, so a downstream model is built on the sequence representations of multiple short genomic sequences yielded from the pre-trained model’s encoder to complete the corresponding downstream task, still using the DNA sequence and cell-type specific chromatin accessibility profiles. The benefits of the pre-training and fine-tuning framework over classical individual task-specific predictive models are two-fold. First, it transfers knowledge from one predictive task to another. Second, it allows comprehensively modelling multiple modalities (Fig.1a,c).

Another innovative design of our EPCOT model is the use of the cell-type specific chromatin accessibility profiles as input to train a cell-type specific model and allow generalizability across cell and tissue types. The pre-training and fine-tuning models are generalizable to new cell and tissue types without requiring additional training. Previous predictive models such as DeepSEA [1] which predict epigenomic features across multiple cell lines from the DNA sequence, cannot be generalized to new cell types. Furthermore, using additional chromatin accessibility data improves the epigenomic feature prediction performance [2] and also benefits downstream tasks such as chromatin contact map prediction.

In addition, EPCOT’s encoder-decoder framework assigns learnable embeddings to the pre-training labels and captures their dependence to predict epigenomic features, as opposed to classical multi-task predictive models where the labels are assumed to be independent. This unique framework allows us to model different TF co-binding activities.

### Cell-type specific epigenomic feature prediction (EFP)

Our cell-type specific epigenomic feature prediction model predicts 245 epigenomic features including 236 TFs and 9 histone modifications. To the best of our knowledge, our EPCOT model predicts the largest number of TFs among existing predictive models [2, 21, 22, 23]. In addition, the majority of these models perform single-task training, which is inefficient to predict multiple TFs or learn the relationships among TFs. In contrast, EPCOT leverages a multi-task training framework capable of predicting 245 epigenomic features in a single model and jointly capturing their dependence.

EPCOT accurately predicted cell-type specific epigenomic features. Here we chose two baseline models scFAN [24] and FactorNet [2] which can be designed to perform multi-task training, and a third baseline using averaged pre-training ChIP-seq signals [25]. We compared them with EPCOT in cross-chromosome and cross-cell type prediction. The four cell lines (K562, MCF-7, GM12878, and HepG2) with the most epigenomic feature profiles from ENCODE were used for training and cross-chromosome testing, and 236 TFs that were available in at least two cell lines were used for training. Five additional cell lines (H1-hESC, A549, HCT116, HeLa-S3, and IMR-90) were used to test in a cross-chromosome and cross-cell type fashion. Overall, EPCOT predicted epigenomic features more accurately than the three baselines (Fig.2a and Extended Data Fig.3b,c,d). Moreover, our model accurately predicted TFs and histone modifications which are important for some downstream tasks such as gene expression prediction and chromatin organization prediction (Fig.2d).

**Fig.2:**
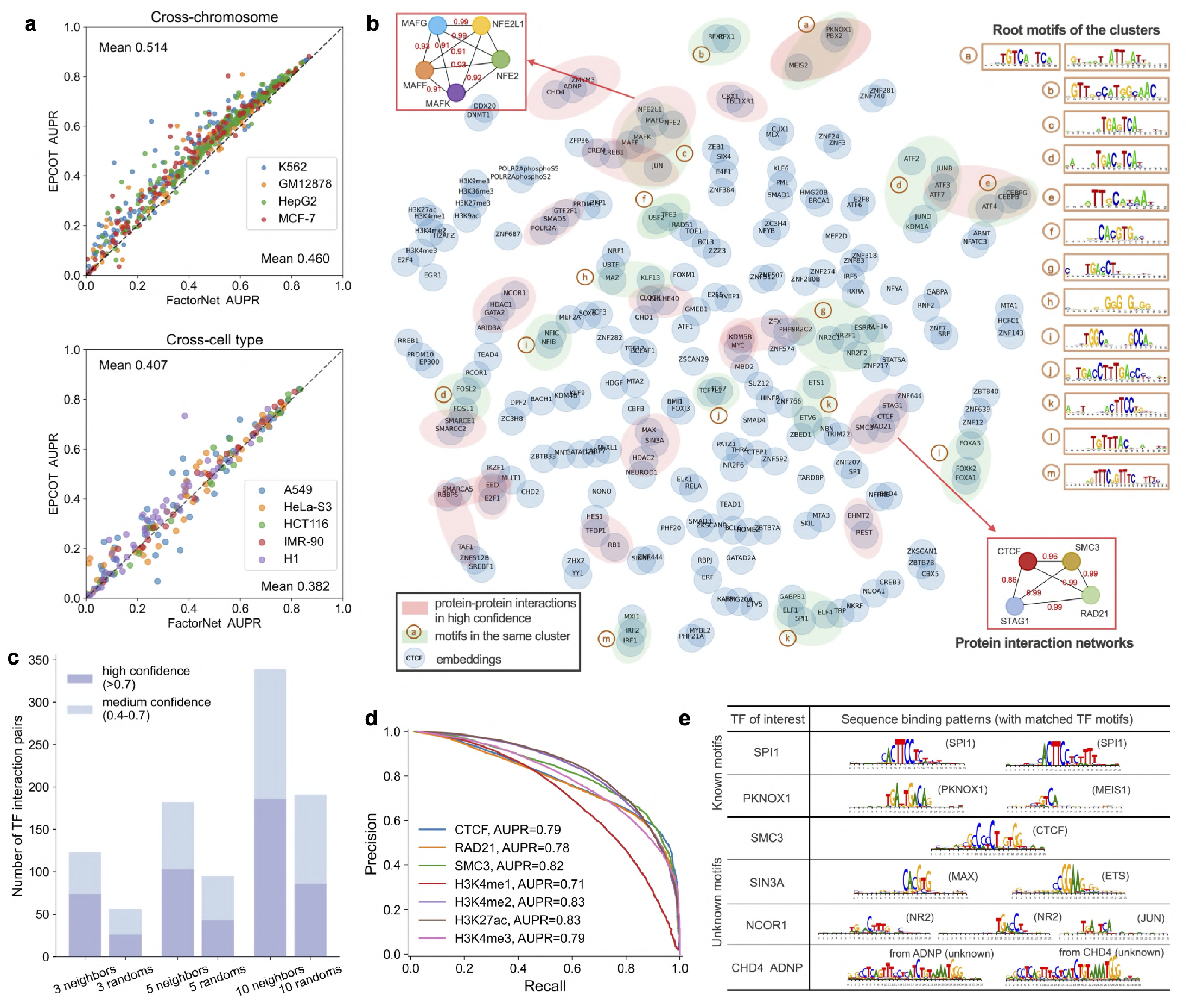
Epigenomic feature prediction and pre-training model interpretation. **a**, EPCOT outperforms FactorNet in cross-chromosome and cross-cell type epigenomic feature prediction. Each dot indicates the AUPR scores of an epigenomic feature predicted by EPCOT and FactorNet. **b**, A *t*-SNE plot of the epigenomic feature embeddings that reflects co-binding patterns. TFs that have high-confidence interactions from STRING [27] or have binding motifs in the same cluster from RSAT [28], are shown in red or green circles, respectively. Additionally, the root motifs of individual binding motif cluster are provided, and two examples of STRING protein interaction networks aligned with TF embeddings are provided, where the scores on edges are the STRING confidence scores of the TF interactions. **c**, The number of TF interaction pairs which are validated by the protein-protein interaction (PPI) dataset STRING [27] at different confidence levels. The nearest neighbors of label embeddings identify more STRING PPIs than the expected number of randomly sampling TF pairs. Three groups are set up where three, five, and ten nearest neighbors are selected for each TF based on the cosine similarity of their label embeddings. **d**, AUPR curves show that EPCOT accurately predicts seven representative epigenomic features which are essential in downstream tasks. **e**, Sequence binding patterns of TFs generated from attribution scores. Some patterns recover known binding motifs. Some patterns match other TFs’ motifs, which reflect co-binding patterns. Some patterns can be unknown binding patterns.

EPCOT’s pre-training model also learns meaningful embeddings for the 245 epigenomic features, which reflects their co-binding patterns. As in natural language processing, if two words frequently appear together in a sentence, the self-attention mechanism captures the dependence between the two words during learning from the corpus. The self-attention mechanism in our pre-training model captures the dependence among these epigenomic features with learnable label embeddings. We hypothesized that if the epigenomic features co-bind frequently, then they can have similar label embeddings. Therefore, we used the protein-protein interaction (PPI) database and known TF binding motifs to validate the hypothesis. First, we leveraged *t*-SNE algorithm [26] to embed the label embeddings into a two-dimension space (Fig.2b). We observed that multiple TFs in local *t*-SNE clusters had high-confidence interactions from STRING database [27]. In addition, since compressing the embeddings into two dimensions might lose important information, next we interpreted their original 512-dimension embedding. We identified the nearest neighbors of each epigenomic feature based on the cosine distance between their embeddings, and expected that the nearest neighbors could be validated by the confidence scores of PPIs from the STRING database. In particular, we set three groups which included three, five, and ten nearest neighbors, respectively, and then counted the number of neighbor pairs that had high and medium confidence scores, respectively (Fig.2c). We observed that nearest neighbors of TF label embeddings identified more STRING PPIs than TF pairs randomly sampled, and 155 unique pairs of neighbors had medium or high confidence interactions in the five-nearest neighbor experiment (Supplementary Table 2). Therefore, the label embeddings of epigenomic features reflected their interaction patterns.

To test if the TFs that have similar label embeddings also have similar binding motifs, we used RSAT clustering results of JASPAR motifs [28, 29]. From the previous *t*-SNE plot, fourteen different TF motif clusters were identified (excluding some motif clusters which contain only one TF such as CTCF and REST). In addition, with the same five nearest neighbors in the higher dimension, we identified 63 pairs of embedding neighbors whose binding motifs were in the same motif cluster (Supplementary Table 1), which reflected that some TFs had similar label embeddings due to their similar binding motifs.

Furthermore, sequence patterns generated from attribution scores reflect TF binding patterns including binding motifs and co-binding patterns. The attribution scores of genomic sequences toward TF binding reflect the important regions that contribute to TF binding prediction, which can be used to generate sequence binding patterns using TF-MoDISco [30], and these sequence patterns are able to recover TFs’ known binding motifs or reflect other sequence binding patterns such as co-binding patterns [31, 32]. From sequence patterns generated for each TF using EPCOT, we first observed that sequence patterns of some TFs recovered their known binding motifs, for example SPI1 (Fig.2e). In addition to known binding motifs, co-binding patterns were also reflected. For example, a known binding motif and a MEIS1 motif were detected in PKNOX1’s sequence patterns, which was supported by a high-confidence interaction score between PKNOX1 and MEIS1 in STRING. Furthermore, for TFs with unknown binding motifs, their sequence patterns could be motifs of other TFs that they have interactions with. For example, the sequence patterns of NCOR1 whose motifs are not listed on JASPAR, included two NR2 family motifs and one JUN motif. In the STRING database, NCOR1 had high-confidence interactions with JUN and NR2 family TFs such as HNF4A, PPARA, and NR2F1. Furthermore, unknown sequence patterns were generated for some TFs. For example, a common but unknown sequence pattern was generated for both ADNP and CHD4, and ADNP interacts with CHD4 to form a complex [33]. Additionally, we observed that CTCF motifs frequently appeared in the sequence patterns of other TFs such as SMC3 and ZNF143 which are shown to interact with CTCF. The generated sequence patterns were available in our GitHub repository, along with their motif comparison results using Tomtom [34] and the STRING scores of the interactions with motif-matched TFs.

### Cell-type specific gene expression prediction (GEP)

EPCOT accurately predicts gene expression including CAGE-seq and RNA-seq in the downstream tasks, and characterizes the relationships between epigenomic features and gene expression. Here we build two downstream models. One is to predict RNA-seq gene expression from genomic regions centered at each gene’s transcription start site (TSS) which is consistent with previous predictive models [5, 6]. The other one is to predict CAGE-seq in a large genomic region depending on the resolution of CAGE-seq to be predicted. Both models take the inputs of DNA sequence and chromatin accessibility data.

In RNA-seq GEP, there exist two tasks, namely binary gene expression classification and gene expression value regression for each protein-coding gene. Since gene expression can be predicted from histone marks, computational models [5, 6, 35] were proposed to predict RNA-seq gene expression from several histone mark profiles within cell types (i,e., using the cell-type specific histone marks to predict gene expression in the same cell type). Compared to these models, EPCOT did not use histone modification profiles as inputs and achieved comparable performances in within-cell type prediction. We first compared EPCOT with GC-MERGE [35] which leveraged six histone marks and chromatin contact maps to predict RNA-seq gene expression in both classification and regression tasks. We observed that EPCOT slightly outperformed GC-MERGE in the two pre-training cell lines (GM12878 and K562) in both tasks, and achieved comparable performances in HUVEC (Extended Data Fig.4a).

EPCOT accurately predicts gene expression for new cell types. In this task, EPCOT was trained on four cell types (H1, A549, GM12878, and HeLa-S3) and tested on four different cell types (K562, Human Mammary Epithelial Cells (HMEC), Human Skeletal Muscle Myoblasts (HSMM), and HepG2). Here, we compared our cross-cell type prediction results with the within-cell type prediction results from DeepChrome [6] and AttentiveChrome [5] which used five histone marks to perform gene expression classification. We observed that EPCOT which used less data, achieved comparable cross-cell type prediction performance with Deepchrome and AttentiveChrome’s within-cell type prediction performance (Fig.3c and Extended Data Fig.4c,d).

**Fig.3:**
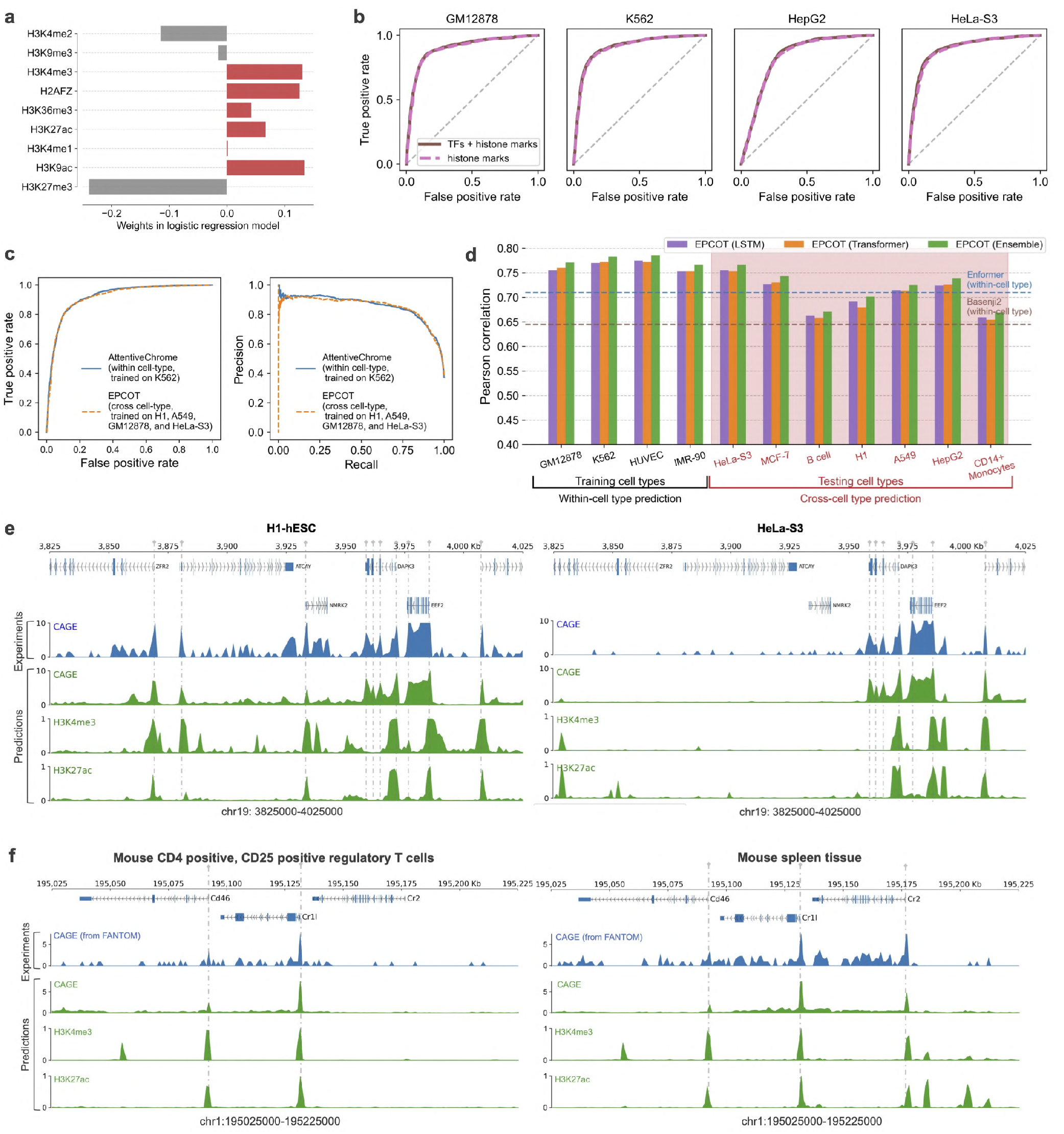
Performance of EPCOT in cell-type specific GEP tasks. **a**, Weights in the logistic regression (LR) GEP model (which uses the predicted values of nine histone marks from the pre-training model) reflect their contributions to GEP. **b**, ROC curves of two LR models (one uses all the 245 predicted epigenomic features and the other one uses nine predicted histone marks only) indicate that histone marks are sufficient to predict gene expression. **c**, EPCOT accurately predicts cross-cell type RNA-seq gene expression which is comparable to AttentiveChrome’s within-cell type prediction in K562. The comparisons of other cell types are provided in Extended Data Fig.4c. **d**, Within-cell type and cross-cell type prediction performance of EPCOT in 1kb-resolution CAGE-seq prediction. EPCOT is trained and tested on four cell types shown as the ‘training cell types’. Cross-cell type evaluation is performed on the remaining seven cell types shown as ‘testing cell types’. Pearson correlation of CAGE-seq (log(1+*x*) transformed) is calculated across all genomic bins in the testing genomic regions. The average Pearson correlation scores reported by Enformer and Basenji2 are shown as two dashed lines. **e**, Predicted CAGE-seq for two new cell lines (H1-hESC and HeLa-S3) at a 200kb region from the EPCOT model trained on GM12878, K562, HUVEC, and IMR-90. The CAGE-seq peaks at gene TSS are predicted and also aligned with predicted H3K4me3 and H3K27ac peaks from the pre-training model. **f**, Cross-species prediction of CAGE-seq and histone marks on mouse cell and tissue types. The CAGE-seq peaks at gene TSS are predicted by the model trained on human cell types, which agree with predicted H3K4me3 and H3K27ac peaks from the pre-training model.

Furthermore, the predicted epigenomic features from the pre-training model further predict gene expression simply through logistic regression (LR) models, which characterize the effects of epigenomic features on GEP. We built two LR models to predict gene expression from the predicted values of all the epigenomic features or nine histone marks from the pre-training model, respectively. We first observed that histone marks were sufficient to predict gene expression since additional TF information did not increase the prediction accuracy (Fig.3b and Extended Data Fig.4b). Furthermore, the weights in the LR model characterized the contributions of histone marks in GEP (Fig.3a). We observed that histone marks related to gene activation all received positive weights, whereas gene repression histones received negative weights, and H3K4me1 which is associated with enhancer regions received a weight close to 0, and the negative weight of H3K4me2 may be due to its presence on poised promoters [36].

In the CAGE-seq GEP task, we compared EPCOT with GraphReg [7] in 5kb-resolution prediction using the same cross-validation strategy to obtain global results. EPCOT significantly outperformed Seq-CNN [7] and Seq-GraphReg [7] which utilized cell-type specific DNase-seq, H3K4me3, and H3K27ac as pre-training and predicted CAGE-seq from DNA sequence either using chromatin contact maps or not (Extended Data Fig.4e). EPCOT also achieved comparable performances with Epi-CNN [7] and Epi-GraphReg [7] which predicted CAGE-seq from cell-type specific DNase-seq, H3K4me3, and H3K27ac tracks either using chromatin contact maps or not.

Furthermore, EPCOT predicts cell-type specific CAGE-seq at 1kb resolution for new cell types. We performed within-cell type prediction on the four training cell types and performed cross-cell type prediction on seven additional cell types (Fig.3d). We separately trained two types of downstream models with two different neural networks, LSTM and Transformer. The two downstream models had comparable performance. Additionally, we tested the ensemble of the Transformer and LSTM models which took an average of the outputs from the two models, and the ensemble model achieved slightly better performance. Furthermore, EPCOT learned cell-type specific CAGE-seq information associated with cell-type specific epigenomic features in the cross-cell type prediction (Fig.3e). To demonstrate the cross-cell type prediction accuracy, we simply compared EPCOT with the average Pearson correlation reported in Enformer [3] and Basenji2 [37] (two dashed lines in Fig.3d). These two models have different training frameworks and different scopes from us. Enformer and Basenji2 perform multi-task prediction and predict CAGE-seq profiles in multiple cell types from the DNA sequence, and cannot be generalized to new cell types. By contrast, EPCOT performs single-task training where the input chromatin accessibility data and CAGE-seq are in the same cell type. EPCOT predicts CAGE-seq for new, unseen cell types if their chromatin accessibility profiles are available. Furthermore, we compared EPCOT’s cross-cell type prediction with prediction using averaged CAGE-seq signal of other cell types, and EPCOT achieved higher correlation on genomic bins associated with gene’s TSS for five testing cell types, and comparable correlation scores for two cell types (Extended Data Fig.4f).

In addition, EPCOT accurately predicts cross-species CAGE-seq. The model trained on human cell types was transferred to mouse cell and tissue types, and predicted differential CAGE-seq peaks in different cell and tissue types (Fig.3f and Extended Data Fig.5).

### High-resolution chromatin organization prediction (COP)

EPCOT predicts high-resolution Hi-C, Micro-C, and ChIA-PET contact maps as one of the downstream tasks (Fig.4a). The downstream models consist of a trunk and a prediction head, the trunk pools and updates the high-level sequence representations from low-level sequence representations output from the pre-training model’s encoder. The trunk contains either LSTM layers or Transformer encoder layers to learn the interactions between different genomic locations. The prediction head is similar to Akita [8], which transforms representations on 1D sequence into representations on 2D contact matrices and predicts the contact maps (Extended Data Fig.2b).

**Fig.4:**
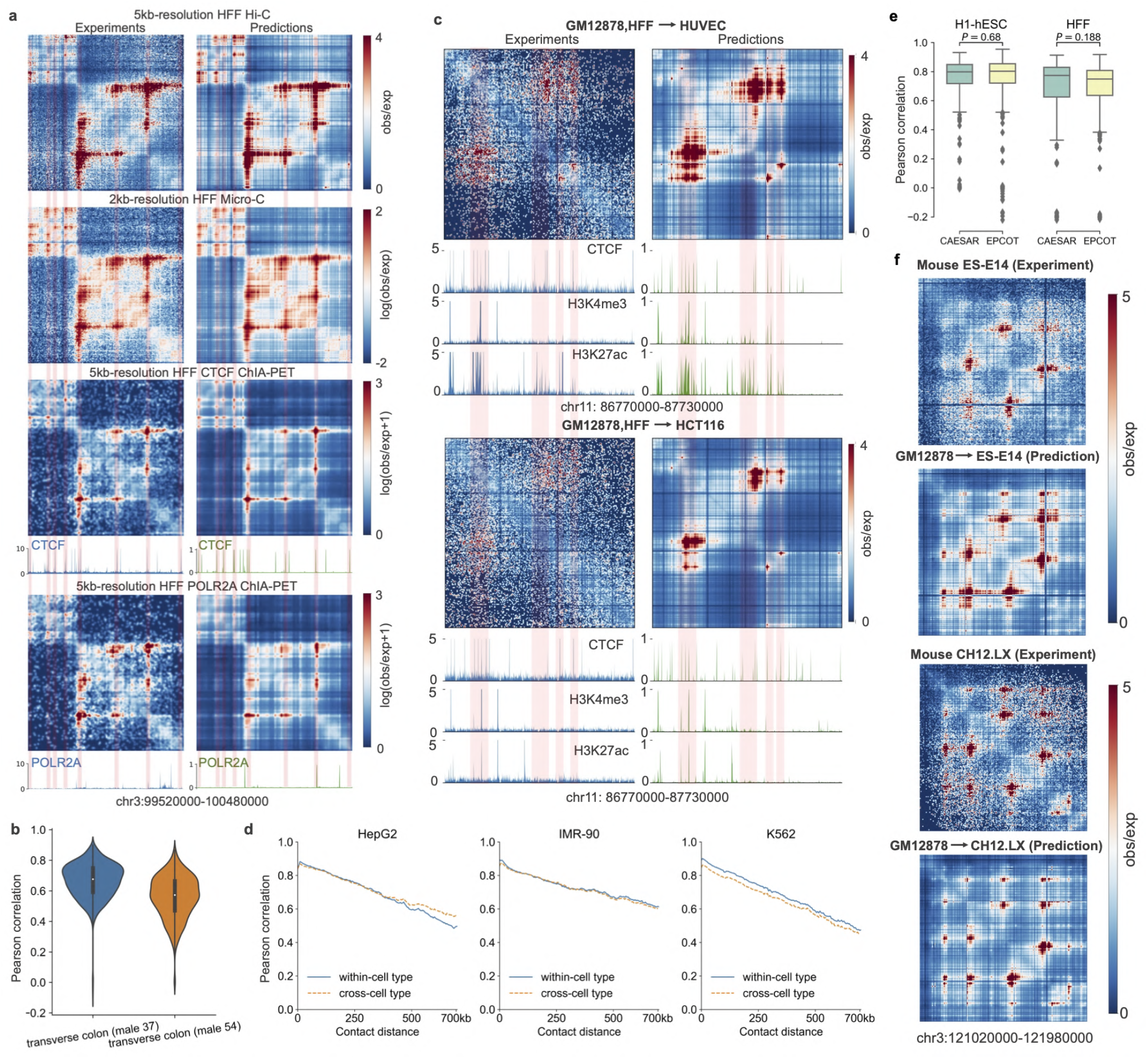
EPCOT accurately predicts high-resolution chromatin contact maps. **a**, EPCOT accurately predicts different types of chromatin contact maps. In an example region (chr3: 99,520,000-100,480,000), predicted HFF Hi-C and Micro-C contact maps, and CTCF and POLR2A ChIA-PET contact maps with the predicted CTCF and POLR2A binding activities are shown on the right. The experiment contact maps and CTCF tracks are shown on the left. The predicted Hi-C, Micro-C and ChIA-PET contact maps agree with the observed contact maps, and the salient regions associate with the CTCF binding sites. **b**, EPCOT predicts Hi-C contact maps for GTEx tissues. Pearson correlation of 1Mb regions in test chromosomes is provided. The cross-donor prediction results are provided in Extended Data Fig.7. **c**, Cross-cell type prediction on two cell lines whose available contact maps have low read depth. The tracks and predicted binding activities of CTCF, H3K4me3, and H3K27ac are provided. The difference in salient regions in the two cell lines is associated with the difference in H3K27ac. **d**, EPCOT predicts cross-cell type Hi-C contact maps, the distance-stratified Pearson correlation in both cross-cell type and within-cell type prediction is shown. **e**, EPCOT achieves comparable performance with CAESAR in 1kb-resolution Micro-C contact map prediction, indicated by the insignificant *p*-values from two-sided Student’s *t*-tests. Pearson correlation is calculated for each 500kb genomic region on test chromosomes. **f**, The mouse Hi-C contact maps (ES-E14 on top, CH12.LX on bottom) predicted by EPCOT trained on human GM12878. The experiment and predicted contact maps in an example region of two mouse cell types are shown, and the differential patterns between these contact maps are predicted.

In Hi-C COP, we predicted the upper triangle of OE-normalized contact matrices in 5kb resolution within 1Mb genomic regions. EPCOT accurately predicted the high-resolution Hi-C contact maps in six cell lines and two GTEx tissue donors (Fig.4b and Supplementary Table 3). We included two baselines, namely Epiphany [38] and DeepC [9] which predict 5kb-resolution Hi-C contact maps. Epiphany predicted OE-normalized Hi-C by using five epigenomic feature tracks, and DeepC predicted percentile-normalized Hi-C from the DNA sequence which also utilized an epigenomic-feature supervised pre-training model. In addition, the two models used 1Mb regions to predict the diagonal only, which required more computational resource than predicting the upper triangular contact matrix of each 1Mb region. The prediction performance reported in the two works and the performance of EPCOT were summarized in Supplementary Table 3. EPCOT achieved higher Pearson correlation and Spearman correlation scores in the same predicted cell lines. Additionally, between the two types of neural networks, LSTM and Transformer in EPCOT downstream model’s trunk, Transformer uniformly achieved better performance than LSTM.

Furthermore, EPCOT also predicted cross-cell type and cross-tissue type Hi-C contact maps. Similar to HiC-Reg [39], an ensemble of two models which trained on the cell lines with the most read counts (GM12878 and HFF), were leveraged to predict four cell lines, namely HepG2, IMR-90, K562, and H1. We observed that EPCOT achieves comparable performance between within-cell type prediction and cross-cell type prediction (Fig.4d and Extended Data Fig.6a). Furthermore, the cross-cell type prediction was compared to Hi-C prediction by averaging the two training cell types’ Hi-C contact maps, and EPCOT achieved better performance in all the cell types (Extended Data Fig.6b). Hi-C contact maps of two cell lines with low read depth were also predicted, the cell-type specific salient regions in contact maps associated with cell-type specific epigenomic features were predicted by EPCOT (Fig.4c). In cross-tissue type prediction, the model trained on traverse colon tissue was used to predict contact maps for two different human tissues (i.e., stomach and tibial nerve), and characterized differential chromatin contacts at the genomic regions with differential chromatin accessibility in the two tissues (Extended Data Fig.7c).

EPCOT also accurately predicted Micro-C contact maps. Here two baselines that predicted high-resolution Micro-C contact maps, Akita [8] and CAESAR [10] were chosen, and Micro-C contact maps in two cell lines H1 and HFF were predicted. The first baseline AKITA predicted Micro-C contact maps in 2,048bp resolution from the DNA sequence, so to be consistent with Akita, we predicted the upper triangle of 2kb-resolution Micro-C contact maps in 1Mb genomic regions. Pearson and Spearman correlation between predicted and target log(observed/expected) contact values were calculated for each bin pair in every region of the test set. EPCOT achieved significantly higher Spearman correlation scores than Akita (Supplementary Table 3), and Akita reported a genome-wide Pearson correlation score 0.61 which was also lower than EPCOT. EPCOT achieved Pearson correlation and Spearman correlation higher than 0.78 in H1 and HFF cell lines. Furthermore, we compared EPCOT with CAESAR in 1kb-resolution Micro-C contact map prediction over 500kb genomic regions. CAESAR utilized much more data than EPCOT, including Hi-C contact maps and six epigenomic feature profiles, to predict Micro-C contact maps, but EPCOT achieved similar performance to CAESAR by only using reference genomic sequence and DNase-seq (Fig.4e), which indicated that genomic sequence and DNase-seq are sufficient to accurately predict high-resolution Micro-C contact maps.

EPCOT also predicts cross-cell and tissue type Micro-C. The model trained on HFF cell line was transferred to predict six cell types and three GTEx tissue donors (Extended Data Fig.6c and Supplementary Figure 2a). Among these cell and tissue types, only H1 had experiment Micro-C contact map. Therefore, in the comparisons between EPCOT’s cross-cell type prediction and H1’s Micro-C prediction using HFF’s contact map, we observed that EPCOT achieved higher average correlation scores (Extended Data Fig.6d). For other cell and tissue types, we compared the predicted Micro-C contact maps with their available experiment Hi-C and observed that the predicted Micro-C not only recovered the Hi-C contact patterns but also characterized fine-scale chromatin interactions (Supplementary Figure 2b).

In addition to comprehensive chromatin organization, EPCOT also predicted chromatin interactions specific to proteins such as ChIA-PET contact maps. The same model architecture to predict 5kb-resolution Hi-C was leveraged here to predict CTCF and POLR2A ChIA-PET in three cell lines. EPCOT accurately predicted ChIA-PET in both within-cell type and cross-cell type prediction (Fig.4a and Extended Data Fig.6f,g). The ChIA-PET contact maps were sparser than Hi-C, but EPCOT predicted the sparse salient regions well in both CTCF and POLR2A ChIA-PET which were also consistent with the predicted binding activities from the pre-training model.

Furthermore, EPCOT predicted cross-species Hi-C contact maps for mouse cell types (Fig.4f). The model trained on human GM12878 was used to predict two mouse cell lines (i.e., ES-E14 and CH12.LX). For both mouse cell lines, Pearson correlation with measured contact maps in this cross-species prediction setup was slightly lower than that in within-cell type prediction (Extended Data Fig.8b,c). EPCOT’s pre-training model also predicted cross-species epigenomic feature profiles. We observed that the salient regions in the contact matrices and their associated epigenomic feature peaks were recovered by EPCOT (Extended Data Fig.8d).

### Enhancer activity prediction (EAP)

EPCOT predicts active enhancers captured by STARR-seq. The active enhancers are predicted from the candidate enhancer set on ENCODE [40] with specific downstream models. Here we focused on the five human cell lines whose STARR-seq profiles were available on ENCODE. STARR-seq measures enhancer activity across the whole genome [41], which allows us to generate the positives (active enhancers) and negatives (inactive enhancers) for model training. To generate the positive samples, we used a similar strategy in the work [12] which combined matched filter scores of five histone marks and DHS to predict Drosophila enhancer activity through a linear SVM model (referred to as “matched-filter model”) — if a candidate enhancer overlapped with STARR-seq and H3K27ac peaks, then we identified it as an active enhancer in high confidence, whereas if a candidate enhancer did not overlap with STARR-seq peaks and was not identified as an active enhancer from EnhancerAtlas [42] either, then we took it as a confident inactive enhancer.

EPCOT accurately predicts cell-type specific enhancer activity. To compare with the matched-filter model, we randomly selected ten times more negatives than positives in the testing data, and EPCOT achieved AUPR scores greater than 0.85 (Extended Data Fig.10a) in all of the five cell lines, whereas the matched-filter model reported AUPR 0.66 in the same positive to negative ratio. Furthermore, EPCOT also accurately predicts active enhancers for unseen cell types. Here we trained EPCOT on HCT116, K562, and MCF-7 cell lines, and tested EPCOT on HepG2 and A549 cell lines. Then we compared the performance of this cross-cell type prediction with the performance of within-cell type prediction using all the positives and negatives in the testing data (Fig.5a). Since the testing data were highly imbalanced (Supplementary Table 4), we focused on the PR curves and observed that the PR curves from cross-cell type prediction were slightly lower than those from within-cell type prediction.

**Fig.5:**
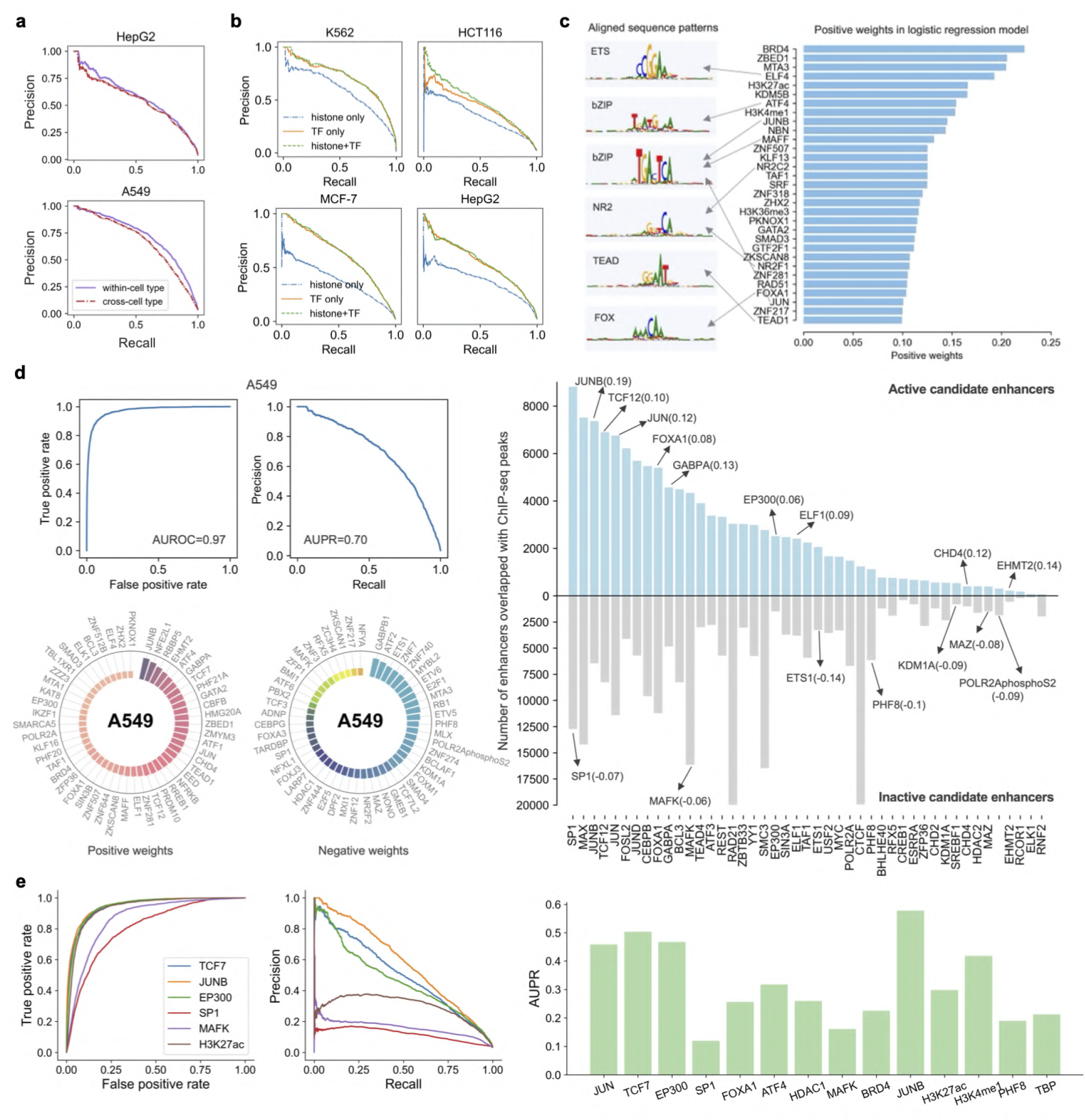
EPCOT accurately predicts enhancer activity and characterizes the effects of epigenomic features to-wards EAP. **a**, The AUPR curves of within-cell type and cross-cell type EAP in two cell lines. EPCOT accurately predicts enhancer activity for new cell types, and the cross-cell type prediction model is trained on K562, HCT116, and MCF-7. **b**, The AUPR curves in EAP using *L*_1_-regularized logistic regression (LR) with the inputs of the 245 predicted epigenomic features, 236 TFs, or nine histone marks, respectively. TFs are more predictive than histone marks in enhancer activity prediction. **c**, The top 30 largest positive negative weights for epigenomic features in the 245-epigenome LR model in all five cell lines. The weights characterize their general effects on enhancer activity. Some of these epigenomic features have motifs agreed with learned sequence patterns from the active enhancer sequences. **d**, Analyzing cell-type specific relationships between enhancer activity and TFs in A549. A549 is not a pre-training cell line whose epigenomic feature profiles are not used in training. (Top left panel) AUROC and AUPR curves of training LR model to predict enhancer activity. (Bottom left panel) 50 TFs with largest positive and negative weights in the LR model. (Right panel) Number of TF ChIP-seq peaks overlapped with active and inactive enhancer regions, and TFs with large weights in the LR model are highlighted. **e**, LR prediction performance using single epigneomic features in A549. AUPR scores of fourteen TFs that receive large positive or negative LR weights are provided. The ROC and PR curves of the 6 representative TFs are shown.

EPCOT also identifies cell-type specific sequence patterns in active enhancer regions (Extended Data Fig.11). The sequence patterns are generated from the attribution scores of DNA sequence toward enhancer activity in an investigated cell line using TF-MoDISco [30]. We observed that the generated sequence patterns matched with TF motifs such as CREB, JUN, YY, ETS, and IRF, which related to enhancer activity and were enriched in enhancers [43].

Similar to gene expression prediction, the enhancer activity can also be predicted by a logistic regression (LR) model from the epigenomic features predicted by our pre-training model. However, different from gene expression prediction, we found that TFs were more predictive than histone marks in enhancer activity prediction, which was also concluded by Dogan et al. [44]. The LR models with all epigenomic features or TFs significantly outperformed the LR model with histone marks only (Fig.5b). Furthermore, the general impacts of epigenomic features on EAP in five cell lines are characterized by the weights in the LR model. First, the weights for five related histone marks in the LR model agreed with those in linear SVM used in the matched-filter model (Extended Data Fig.10b). Next, multiple epigenomic features that received large positive weights were shown to be associated with active enhancers in the literature (Fig.5c), such as BRD4, MTA3, RAD51, H3K27ac, and H3K4me1 [45, 46, 47, 48], and some of these TFs also have motifs similar to learned sequence patterns, such as ATF4, ELF4, and JUNB. Additionally, some TFs that bind to promoter regions such as PHF8, TBP, and SP1, and some TFs related to repression activity such as HDAC1, received large negative weights.

Furthermore, the importance of TFs to EAP is captured in a cell-type specific manner. We found that the LR models separately trained for individual cell lines outperformed the LR model trained on all five cell lines (Extended Data Fig.10d), which indicated that the effects of some epigenomic features to enhancer activity can be cell-type specific. To analyze the cell-type specific relationship between TFs and enhancer activity, first the high prediction accuracy indicates that the weights in LR capture the contributions of TFs to EAP (Fig.5b,d). Although TFs with large weights in the LR model can be different in different cell lines, we observed that some TFs such as ELF4, BRD4, ATF4, and RBBP5, received large positive weights, and PHF8 and HDAC received negative weights in the three investigated cell lines, MCF-7, A549, and K562 (Fig.5d and Extended Data Fig.10e). Furthermore, we found that some TFs received large positive weights due to their frequent binding on active enhancer regions (Fig.5d right panel), such as JUNB, TCF12, and JUN. The LR weights of a large number of epigenomic features may not reflect their true contributions to enhancer activity prediction due to their correlations such as co-linearity. Therefore, enhancer activity prediction using single epigenomic features was also performed (Fig.5e). We observed that some TFs with negative weights such as SP1 and MAFK predict enhancer activity less accurately than some TFs with large positive values such as JUN and TCF7. The majority of the cell types only have a limited number of TF profiles, which prevents us from capturing cell-type specific relationships between TFs and enhancer activity, but EPCOT predicts 236 TFs for these cell types and quantifies their contributions, which helps us understand cell-type specific impacts of TFs on enhancer activity.

### Generic sequence representations to predict multiple modalities

The pre-training model’s encoder, supervised by epigenomic features, learns sequence representations from the input DNA sequence and chromatin accessibility data, which is then fine-tuned in each of the downstream prediction tasks (i.e., sequence representations are updated during fine-tuning). We have observed that individual sequence representations yielded from the original pre-training model and updated for each of the downstream tasks perform well in the corresponding task. However, one may ask if there exists globally optimized (generic) sequence representations that can be used in all of the prediction tasks. For example, can the pre-training model fine-tuned in the GM12878 COP task be frozen and transferred to the K562 GEP task? These questions are intriguing because we are searching for generic sequence representations that computers understand and use to decode the human genome and complete these genomic prediction tasks.

In order to address these important questions, we first tested if the sequence representations yielded from the original pre-training model (EFP task) could be fixed to predict other tasks. We have observed that the predicted epigenomic features from the pre-training model accurately predicted RNA-seq gene expression and enhancer activity through a simple logistic regression model, which indicated that sequence representations yielded from the frozen pre-training model were able to perform well in the two tasks. However, fine-tuning the parameters in pre-training model’s encoder significantly outperformed fixing these parameters in the Hi-C COP task (Extended Data Fig.9a), which indicated that genomic sequences possibly contained information in addition to epigenomic feature binding that helped Hi-C contact map prediction.

Then we tested if the original pre-training model that was fine-tuned in the Hi-C COP task could produce generic sequence representations. We fine-tuned the original pre-training model in GM12878 Hi-C prediction, since GM12878’s Hi-C data [49] had higher read depth than other cell types, and the sequence representations output from this fine-tuned model were then transferred to train downstream models to predict additional tasks. Overall, the globally optimized sequence representations achieved comparable performance with individual sequence representations updated in each prediction task, and the fine-tuned pre-training model is generalizable to cell types other than GM12878 (Fig.6). Although in some tasks such as Micro-C COP and CAGE-seq GEP, the generic sequence representations achieved worse performance, the performance did not differ too much, which was as expected since individual representations optimized for each specific task and cell type should have the best performance. Therefore, the global sequence representations yielded from the pre-training model that is fine-tuned in the GM12878 Hi-C COP task, can be used to achieve satisfactory performances in all of the prediction tasks for other cell types.

**Fig.6:**
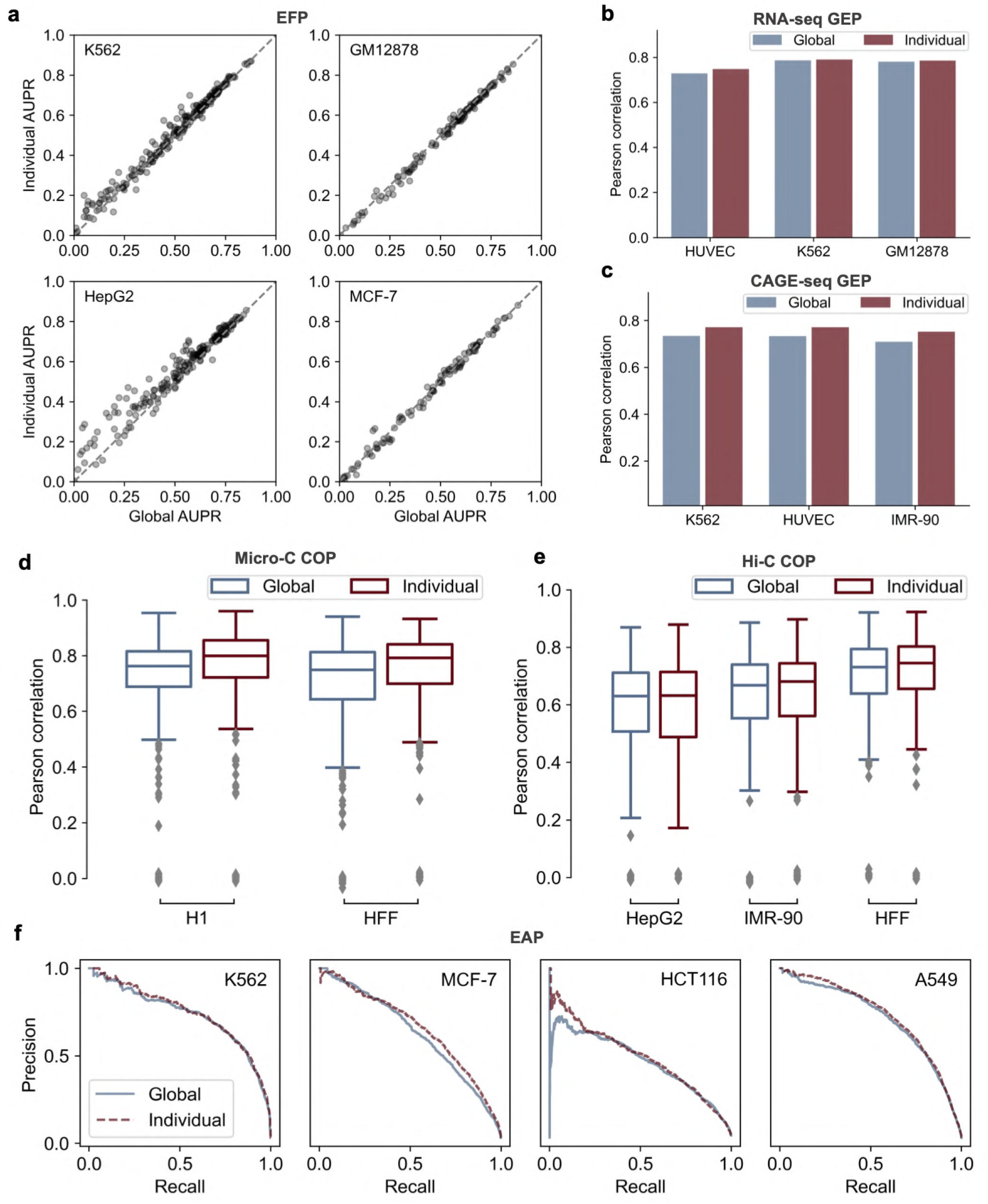
Globally optimized sequence representations perform well in all prediction tasks. ‘Global’ refers to the sequence representations yielded from the original pre-training model (EFP task) that is fine-tuned in GM12878 Hi-C COP task and then fixed in additional prediction tasks including EFP, GEP, COP, and EAP tasks. ‘Individual’ refers to the representations yielded from the original pre-training model that is then fine-tuned and optimized in each of the additional prediction task. Global representations achieve comparable performance with individual representations in EFP (**a**), RNA-seq GEP (**b**), CAGE-seq GEP (**c**), Micro-C COP (**d**), Hi-C COP (**e**), and EAP (**f**) tasks.

## Discussion

EPCOT adopts a novel pre-training and fine-tuning framework to comprehensively predict epigenome, chromatin contact maps, transcription, and enhancer activity. Unlike previous predictive models which predict different modalities separately, EPCOT predicts epigenomic features in the pre-training task and transfers the epigenome-related knowledge to predict other modalities in downstream tasks. In addition, different from previous DNA-sequence based predictive models [1, 8, 3] that make predictions from DNA sequence only, EPCOT takes both DNA sequence and chromatin accessibility data as input, which can predict for unseen cell and tissue types. Moreover, previous self-supervised pre-training works [15] and other models [7, 9] which use supervised pre-training techniques on DNA sequence only, cannot be generalized to new cell and tissue types either. Different from imputation-based models [50, 51, 52], EPCOT predicts for unseen cell types without requiring additional training and predicts many more TF binding profiles, and EPCOT also utilizes DNA sequence information which allows sequence-based analysis. Additionally, EPCOT trained on human cell types can be transferred to mouse species. Furthermore, EPCOT learns global sequence representations which are generalizable among different downstream tasks. The pre-training model which is trained on epigenomic features and then fine-tuned in the GM12878 Hi-C COP task, learns global representations which can be used to accurately predict in other tasks without further fine-tuning. More importantly, this particular fine-tuned pre-training model can be frozen and transferred to complete all of the predictive tasks for new cell types. Therefore, EPCOT’s global representations are different from the sequence representations learned from previous pre-training models [9, 15] that are updated in specific tasks. Its generalizability both across different predictive tasks and across different cell types convinces us that the representations learned by EPCOT are real and reflect underlying biological mechanisms.

Other than superior or comparable predictive performances by only using chromatin accessibility data in different tasks, EPCOT also showed unique methodological advantages. In the EFP task, EPCOT’s pretraining model leverages a multi-task training framework and accumulates training loss only on available epigenomic feature profiles, which allows EPCOT to predict the largest number of TF binding profiles in a single model, in contrast with multiple cell-type specific TF binding prediction models [2, 21, 22, 23]. In the GEP task, EPCOT accurately predicts cell-type specific gene expression, which can be generalized to diverse new cell types. However, multiple predictive models [3, 4, 37, 53] cannot perform cross-cell type prediction. Although some works [5, 6, 7, 35] use cell-type specific histone mark profiles as input, they rarely investigate cross-cell type prediction or only investigate the generalizability across GM12878 and K562. In the COP task, EPCOT accurately predicts cell-type specific comprehensive chromatin contact maps and protein-specific chromatin interactions. Different from the models [8, 9, 11] that predict contact maps from the DNA sequence only, EPCOT utilizes additional cell-type specific chromatin accessibility data, which allows EPCOT to predict cell-type specific chromatin contact maps and requires less cell-type specific data than the models [38, 39] which need additional ChIP-seq data. In the EAP task, EPCOT integrates both TF and histone modification information to predict active enhancers captured by STARR-seq [41]. By contrast, previous enhancer prediction works [12, 13, 44] mainly use histone mark data, and barely utilize TF binding or use a limited number of TFs to predict enhancer activity. Since TFs can be more predictive than histone marks [44], which is also validated by EPCOT, we believe that using additional TF binding information makes EPCOT outperform previous models. Although some models [14, 43] predict enhancer activity from the DNA sequence, their model interpretation mainly focuses on characterizing sequence motif importance in the prediction task. By contrast, EPCOT not only captures sequence patterns in active enhancer regions but also characterizes the contributions of specific TFs (including TFs with unknown motifs) to EAP.

EPCOT is interpretable, which reflects biological insights and facilitates scientific discoveries. First, EPCOT’s pre-training model learns meaningful embeddings and sequence binding patterns of epigenomic features, which reflects their co-binding patterns. By contrast, multiple current TF binding prediction models barely capture the dependence among TFs in the prediction, although some predictive models [54] capture the interactions among TF binding motifs in model interpretation. EPCOT also generates sequence binding patterns for multiple TFs from attribution scores of DNA sequences. Some patterns match motifs of other TFs, and some patterns are unknown binding patterns, which helps researchers investigate TF binding mechanisms.

Second, EPCOT captures cell-type specific effects of epigenomic features on enhancer activity and gene expression. The predicted cell-type specific epigenomic features from the pre-training model further accurately predict gene expression and enhancer activity through logistic regression, which quantifies the contributions of epigenomic features to the prediction. However, previous cell-type specific TF binding prediction models [2, 21, 22] barely perform these downstream analyses. In addition, directly using epigenomic feature profiles to characterize their effects on enhancer activity in a cell-type specific manner, is difficult due to the limited epigenomic feature profiles available for the majority of cell types on ENCODE. Although we only investigate with a logistic regression model, other feature selection methods such as decision tree, can also be leveraged to evaluate the contribution of epigenomic features.

Currently, we leverage 236 TFs, although a small portion of all human’s 1,600 TFs [55], which have already yielded exciting results. When more public ChIP-seq data is available in the future, we expect the pre-training model’s decoder to characterize the dependence among additional TFs. From the technical perspective, we are able to redesign the decoder structure [56] to efficiently scale to thousands of labels (a.k.a. TF binding profiles) without much computational burden. Another future direction is to leverage self-supervised pre-training techniques to replace the supervised pre-training strategy in current EPCOT. The supervised labels (i.e., epigenomic features) used in our pre-training model may restrict the extracted information to these predicted epigenomic features, and other useful information contained in the genomic sequence might be neglected. Considering the current success of self-supervised pre-training in natural language processing and computer vision, a self-supervised pre-training framework to extract massive cell-type specific information across different cell types is promising in genomics.

## Methods

EPCOT leverages a pre-training and fine-tuning framework (Fig.1b). In the pre-training stage, a cell-type specific pre-training model is supervised by epigenomic features, which captures the dependence among epigenomic features and learns epigenomic feature related representations from the input genomic sequence and chromatin accessibility data. In the fine-tuning stage, task-specific downstream models are built, and the pre-training model is transferred and fine-tuned to complete downstream tasks. The detailed structures of the pre-training model and downstream models are discussed in the following sections.

### Pre-training model

EPCOT’s pre-training model takes the inputs of 1.6kb DNA sequence (1kb centeral sequence with 300bp flanking sequences upstream and downstream) and the corresponding cell-type specific chromatin accessibility data to predict epigenomic features. The model structure is similar to Query2Label [20], which assigns learnable embeddings to the labels (i.e., epigenomic features) and consists of encoder and decoder parts. The encoder contains convolutional layers and one Transformer [57]’s encoder layer, which learns a series of sequence representations *h*∈ℝ^*n×d*^ from the inputs *X* where *n* indicates the number of representations and *d* indicates the dimension of the representations,

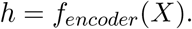

The decoder contains two blocks, and each decoder block consists of a self-attention module, an encoder-decoder attention module (cross-attention module), and position-wise feed-forward networks (FFNs). To capture the dependence among epigenomic features, learnable label embeddings *L*_0_ are first assigned to each of the labels (i.e., epigenomic features) to be predicted, which are then updated through the two decoder blocks.

In the *i*-th decoder block, the updated label embeddings *L*_*i−*1_ from the previous (*i −* 1)-th decoder block are first input to a self-attention module, where a multi-head attention learns the relationships between the labels and updates the label embedding. In this module, the queries, keys, and values of multi-head attention are all from the label embeddings,

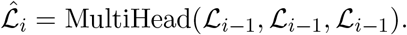

Next, the intermediate label embeddings 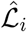 output from the self-attention module with the sequence representations *h* output from the encoder are input to a cross-attention module, where a multi-head attention with keys and values from the sequence representations and queries from label embeddings to select and combine learned representations of interest and update the label embeddings

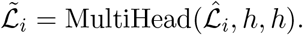

After the two modules, an FFN is applied to the updated label embedding 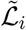

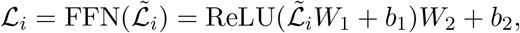

where *W*_1_ and *W*_2_ are two weight matrices, and *b*_1_ and *b*_2_ are bias vectors.

Finally, a label-specific fully connected layer is applied to the final updated label embeddings ℒ_*m*_*∈ℝ*^*N×d*^ through *m* decoder blocks, where *N* represents the number of labels to be predicted and *d* indicates the dimension of the label embedding. The predicted score of *i*-th epigenomic feature *S*_*i*_ is calculated as

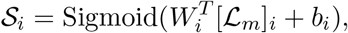

where *W*_*i*_∈*ℝ*^*d*^ is a weight vector corresponding to *i*-th label.

The pre-training model leverages a multi-task training framework, but not all of the 245 target labels are known in the pre-training cell types (Supplementary Figure 1, if an epigenomic feature is available in at least two pre-training cell lines, then it is taken as a predicted label). Therefore, we only calculate the crossentropy loss between the target labels and the predicted scores. In the model training, we first pre-train the convolutional layers on the same target labels, and then train the entire pre-training model by using AdamW optimizer [58] with weight decay of 1*×*10^*−*6^ and an initial learning rate of 5*×*10^*−*4^. The pre-trained encoder is then utilized in the downstream tasks, whose parameters can either be frozen or fine-tuned in the training of downstream models. The hyperparameters of pre-training model are provided in Supplementary Figure 3a.

### Task-specific downstream models

The downstream models are built on the pre-training model’s encoder, which take the inputs of sequence representations *h* output from pre-training model’s encoder. Before feeding the sequence representations into the downstream model, the features *h*∈ℝ^*n×d*^ of each 1kb genomic region are pooled into a 1kb-sequence embedding *ϕ∈ℝ*^*d*^ by using the same attention pooling strategy in Enformer [3], the *j*-th element in the pooled feature vector is calculated as

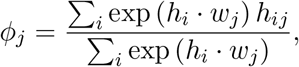

where *w∈ℝ*^*d×d*^ is a learnable weight matrix and *i* is the index of the *n* sequence representations. The downstream models have different architectures for different downstream tasks (Extended Data Fig.2).

### Gene expression prediction (GEP)

In the RNA-seq GEP task, the downstream model takes the input of eleven pooled 1kb-sequence embeddings which represent a 11kb genomic region centered at gene TSS, then we simply apply Bi-LSTM layers or convolutional layers to generate a feature vector which represents the whole input genomic region (Extended Data Fig.2a left). Finally this feature vector goes through a fully connected layer to predict the gene expression values. Considering there are two tasks, binary gene expression classification and gene expression value regression, in the classification task, a Sigmoid function is applied to the outputs of the final fully connected layer and a cross-entropy loss is calculated. By contrast, an MSE loss is calculated in the regression task.

In the gene expression classification task, we also apply logistic regression (LR) to the predicted epigenomic features from the pre-training model. Here the entire pre-training model is frozen and transferred to obtain the predicted values of epigenomic features. Then, we take the maximum predicted values in each 3kb genomic region centered at the gene TSS, which are used to predict gene expression through a *L*_1_-regularized LR model. The weights in LR model quantify the contributions of epigenomic features to GEP.

The downstream model that predicts 1kb-resolution CAGE-seq, takes the inputs of 250kb genomic regions to predict CAGE-seq on the centered 200kb region (Extended Data Fig.2a right). We first apply convolutional layers to the 1kb-sequence embeddings, which learns local relationships among 1kb genomic regions. Then LSTM layers or Transformer’s encoder layers with the same relative positional encoding in Enformer [3] are applied to learn long-range interactions among 1kb genomic regions and update the sequence embeddings. Finally, the updated sequence embeddings are fed into a fully connected layer to predict CAGE-seq tracks. The downstream model architecture and hyperparameters are provided in Supplementary Figure 3b. A simple MSE loss is calculated between the predicted and target CAGE-seq and is optimized by AdamW optimizer with weight decay of 10^*−*6^ and learning rate of 10^*−*4^. In CAGE-seq prediction task, the training batch size is 2 due to the long sequences as input, so BatchNorm [59] used in the model may fail to estimate the population statistics in the inference stage. Therefore, in cross-cell type prediction, we utilize batch statistics in BatchNorm instead of population statistics and set batch size to be 1 to predict one genomic region at each time.

### Chromatin contact map prediction (COP)

The downstream model to predict 5kb-resolution Hi-C contact map takes the inputs of 1Mb genomic regions and predicts the upper-triangle of contact matrices in the centered 960kb, which contains a trunk and a prediction head (Extended Data Fig.2b and Supplementary Figure 3c). In the trunk, the pooled 1kb-sequence embeddings *ϕ* are fed into convolutional layers to learn the local relationships among 1kb genomic regions. Since we now focus on a 5kb resolution, a max-pooling layer is applied to pool five 1kb-sequence embeddings into one 5kb-sequence embedding. Here we use ψ*∈R*^*N×d*′^ to represent the whole feature matrices of the input genomic regions (a stack of 5kb-sequence embeddings), where *N* indicates the number of 5kb genomic regions and *d* indicates the dimension size of 5kb-sequence embedding. Next, we use three Transformer’s encoder layers or LSTM layers to update the embeddings. After learning feature representations for each 5kb genomic region of 1D sequence, to predict the 2D chromatin contact matrices, we transform the updated 2D feature matrices 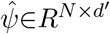 into 3D feature maps Ξ*∈R*^*N×N×d*′^. Each element Ξ_*i,j*_ is a vector related to the chromatin contact between genomic regions *i* and *j*, which is calculated as

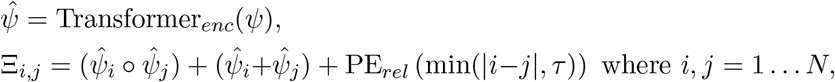

where *i* and *j* index the 5kb genomic regions, and ○ indicates an element-wise multiplication, and PE_*rel*_ represents a relative positional encoding based on the absolute distance between two genomic regions, and *τ* is a threshold to make long-range interactions have the same positional encoding. Then similar to Akita [8], we treat the feature maps Ξ as images and apply several symmetric 2D dilated convolutional layers with skip-connection

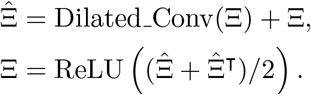

Finally, we apply a fully connected layer to predict the symmetric chromatin contact matrices, and calculate an MSE loss between the predictions and the targets. The downstream model is trained with AdamW optimizer with weight decay of 10^*−*6^ and learning rate of 3*×*10^*−*4^, and we warmup the learning rate in the first training epoch.

The downstream model to predict ChIA-PET contact maps have the same model architecture to the downstream model which predicts Hi-C contact maps. The downstream model to predict 1kb-resolution Microc contact maps also has a similar model architecture, except for a maxpooling layer that pools 1kb-sequence embeddings into 5kb-sequence embeddings.

In this task, the training batch size is 1 because the input sequence is quite long. Since in the training and inference stages, the batch sizes are both 1, similar to the CAGE-seq prediction task, we also use batch statistics in BatchNorm for cross-cell type prediction.

### Enhancer activity prediction (EAP)

The downstream model to predict enhancer activity takes the inputs of 3kb genomic regions (two 1kb bins downstream and upstream the central 1kb enhancer bin are used as flanking regions). Then the pooled sequence embeddings of each 1kb region *ϕ* are fed into convolutional layers and one fully connected layer to predict the enhancer activity score. Similar to the GEP task, enhancer activity can also be predicted by logistic regression using the predicted values of epigenomic features on 1kb enhancer bins, and the parameters of pre-training model are frozen.

## Data availability

All the genomic data used in EPCOT is in the reference genome version hg38. DNase-seq, human CAGE-seq, ChIP-seq and STARR-seq are all downloaded from ENCODE [40]. Mouse CAGE-seq profiles are downloaded from FANTOM [60]. RNA-seq gene expression level data is from REMC database [61]. Hi-C, Micro-C, and ChIA-PET contact maps of cell types are downloaded from 4DN [62], and GTEx tissue Hi-C contact maps are downloaded from ENCODE. Generated TF sequence binding patterns along with their motif comparison results are provided in a web page that is available in our GitHub repository https://github.com/liu-bioinfo-lab/EPCOT.

## Code availability

Source code of EPCOT and trained EPCOT models are available in a GitHub repository https://github.com/liu-bioinfo-lab/EPCOT. We provide codes on the GitHub repository to show how to run EPCOT models on Google Colab notebook to reproduce our experiments and predict multiple modalities for new cell and tissue types if their chromatin accessibility profiles are available.

## Extended Data

**Extended Data Fig.1:**
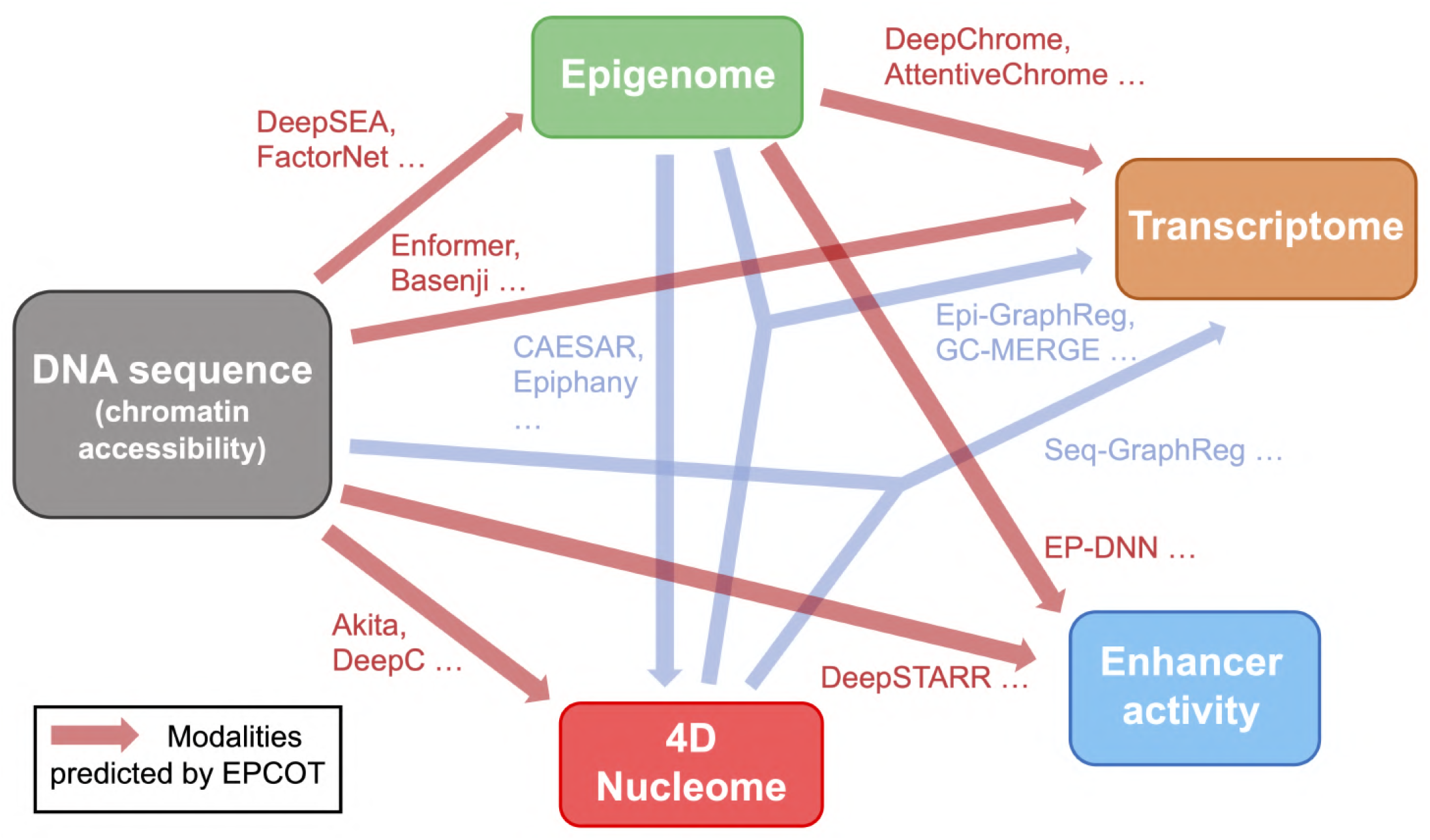
Computational works capturing epigenome, transcriptome, chromatin organization, and enhancer activity.

**Extended Data Fig.2:**
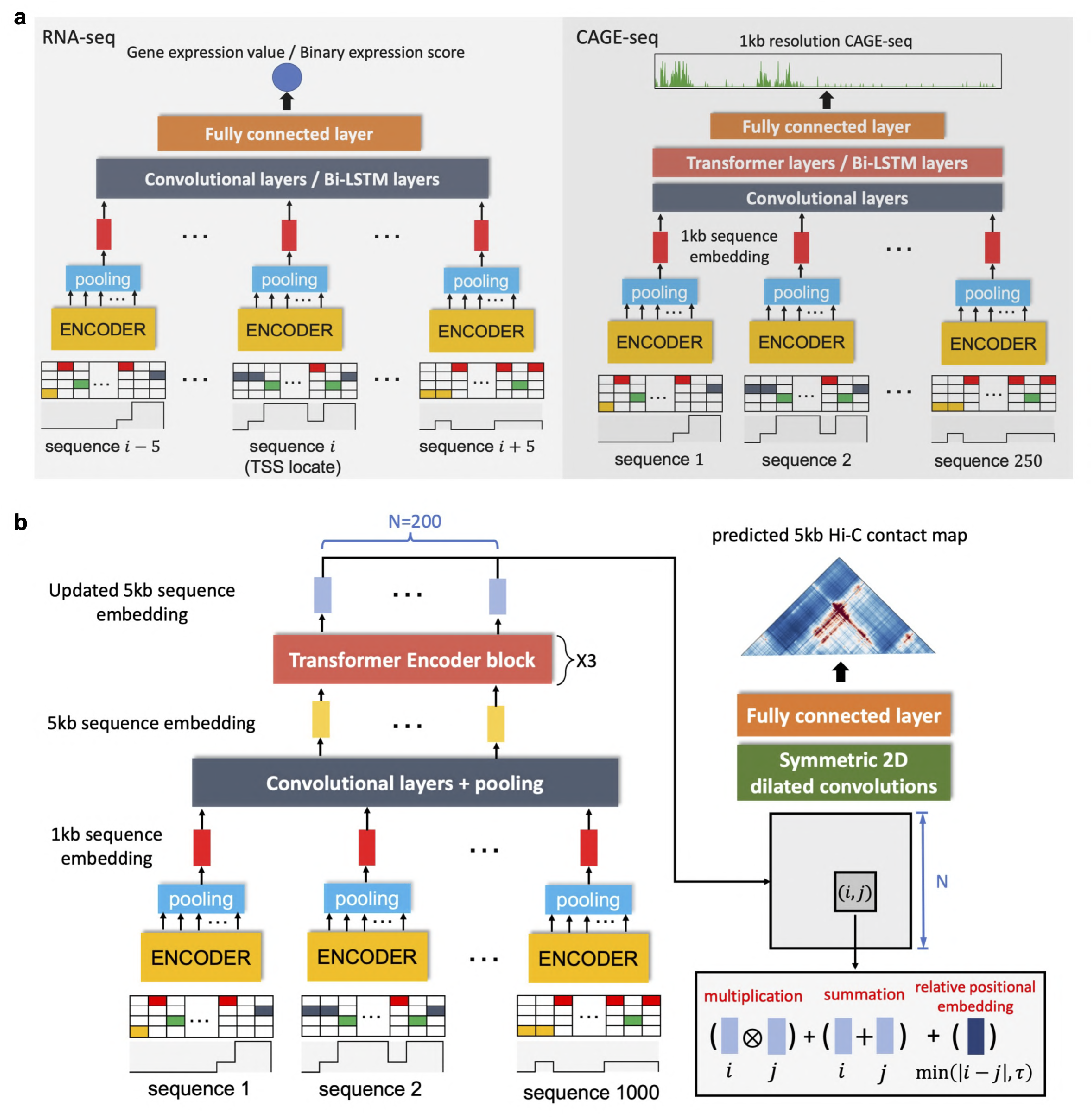
Downstream model architecture. **a)** The downstream models to predict RNA-seq gene expression values and 1kb resolution CAGE-seq tracks, both take the inputs of DNA sequence and chromatin accessibility data. The downstream model to predict RNA-seq uses 11kb genomic regions centered at TSS, and the downstream model to predict 1kb-resolution CAGE-seq uses 250kb genomic regions to predict CAGE-seq on the centered 200kb regions. **b)** The downstream model to predict 5kb-resolution Hi-C contact maps, takes the inputs of 1000 1kb genomic regions. The 1kb-sequence embeddings from pre-training model’s encoder are pooled and updated into 5kb-sequence embeddings which are used to learn the feature representations of the chromatin contacts and predict the upper triangle of the contact matrices.

**Extended Data Fig.3:**
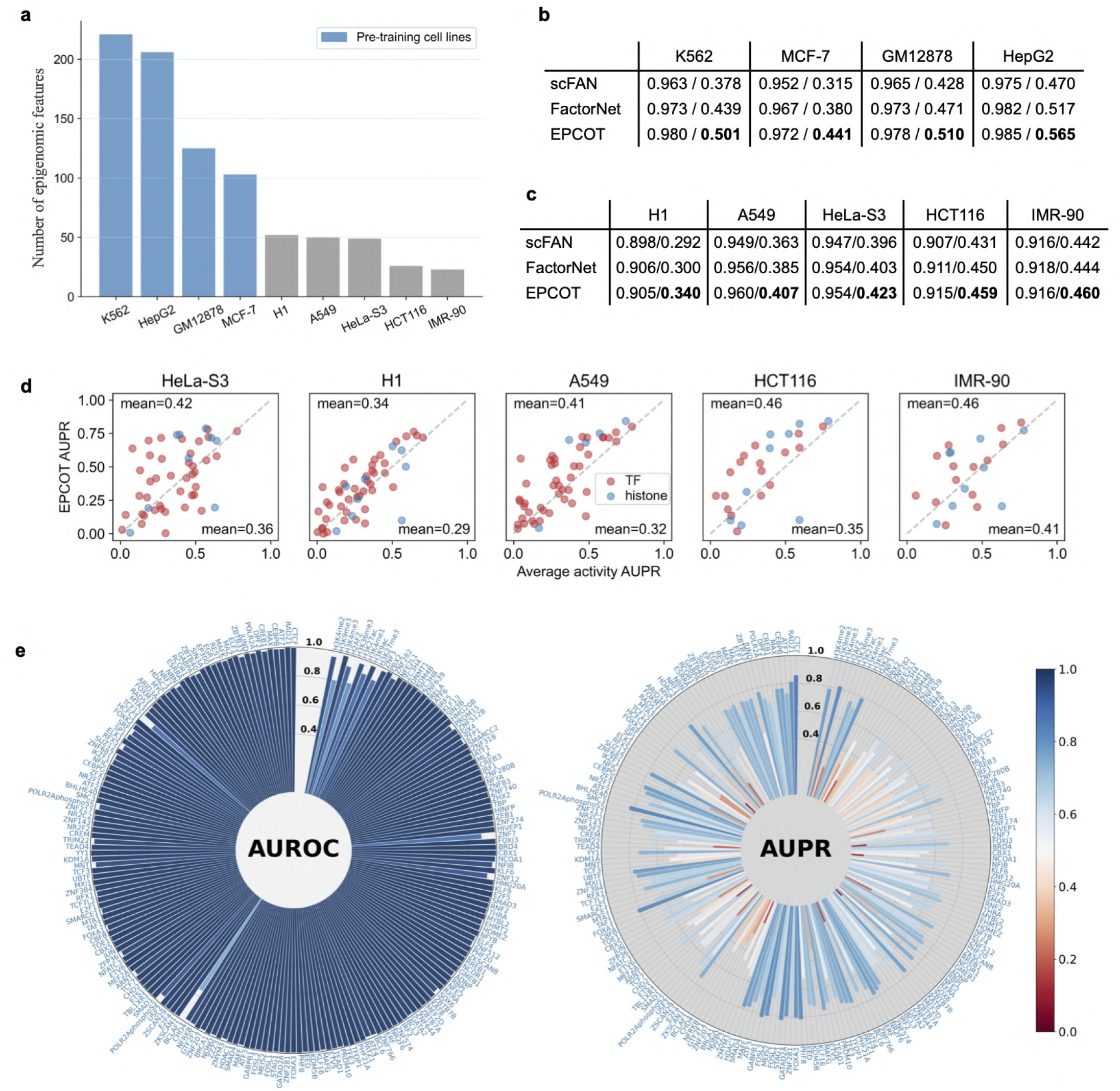
Performance in epigenomic feature prediction. **a)** The number of epigenomic feature profiles collected from ENCODE in nine cell lines. The four cell lines which have the most available profiles are used for pre-training. **b)** Cross-chromosome prediction performance in four pre-training cell lines. The scores (mean AUROC/mean AUPR) of two baselines and EPCOT are provided, and ECPOT outperforms the two baselines. **c)** Cross-chromosome and cross-cell type prediction performance in the remaining cell lines. ECPOT outperforms the two baselines with higher mean AUPR scores. **d)** Comparisons of AUPR scores between EPCOT’s cross-chromosome and cross-cell type prediction and averaging pre-training ChIP-seq signals in five testing cell types. EPCOT achieves higher mean AUPR scores. **e)** AUROC and AUPR scores of 206 epigenomic features in HepG2 cell line. The AUROC scores are generally above 0.9, and majority of epigenomic features achieve AUPR score greater than 0.5.

**Extended Data Fig.4:**
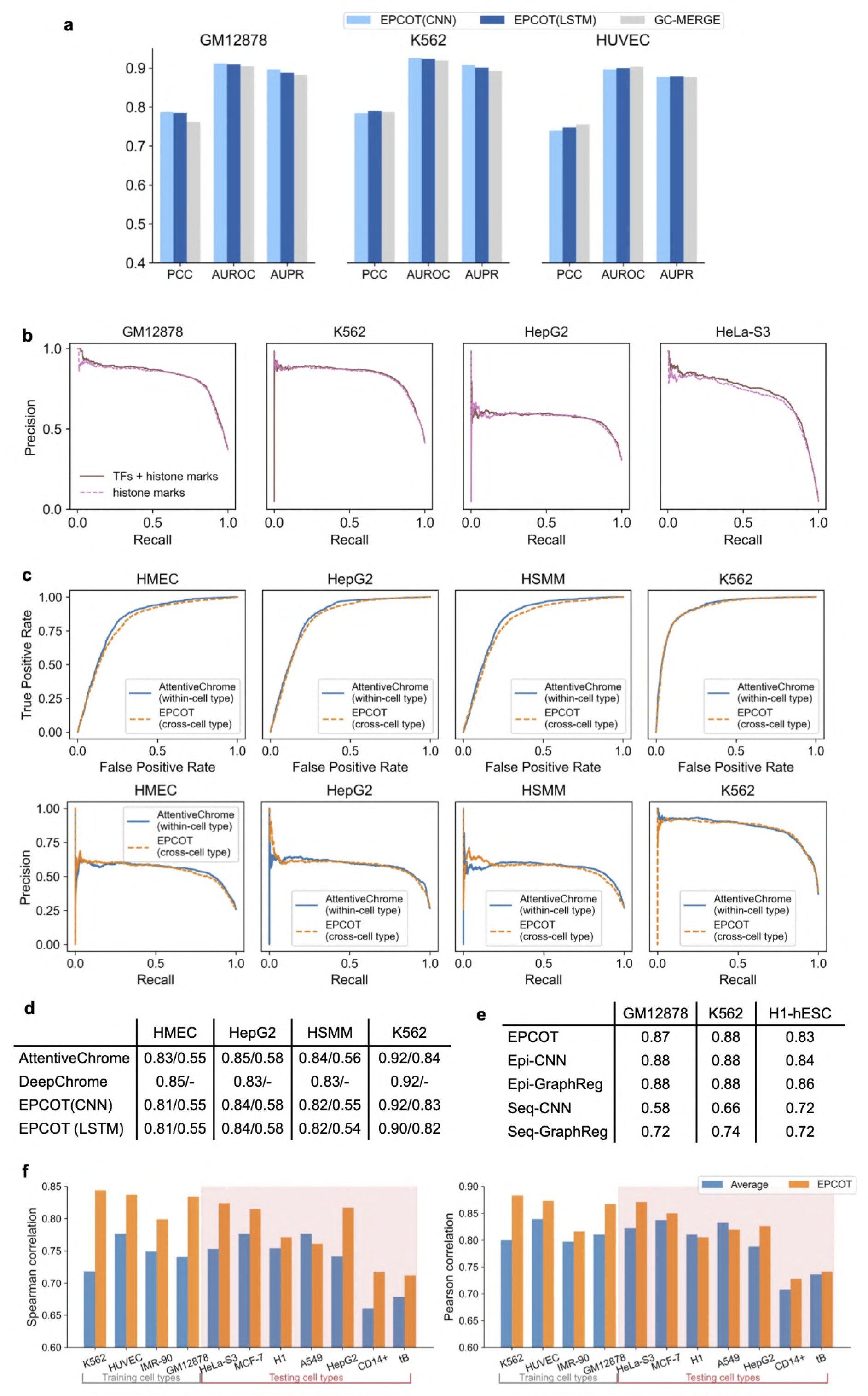
Comparisons in gene expression prediction. **a)** EPCOT (both CNN and LSTM in down-stream models) achieves comparable performance with GC-MERGE in RNA-seq gene expression regression and classification tasks in three cell lines, GM12878, K562, and HUVEC. ‘PCC’ stands for Pearson correlation which is the performance criterion used in the gene expression value regression task, while AUROC and AUPR measure the binary gene expression score classification task. **b)** Comparing performance of using logistic regression to predict gene expression from the predicted scores of all the epigenomic features or histone marks only in four cell lines. The PR curves are shown. **c**,**d)** Comparing EPCOT’s cross-cell type prediction performance with DeepChrome and AttentiveChrome’s within-cell type prediction performance. The ROC and PR curves of EPCOT and AttentiveChrome are shown in **c**, and the AUROC and AUPR scores are provided in **d**. DeepChrome’s AUROC scores are directly from the paper, and the AUROC and AUPR scores of AttentiveChrome are calculated by using Kipoi [63]. EPCOT’s model is trained on four different cell lines: H1, A549, GM12878, and HeLa-S3. **e**) Comparing EPCOT with four models proposed by GraphReg in three cell lines: GM12878, K562, and H1-hESC. Pearson’s correlation across genomic bins associated with gene TSS are calculated, and the scores of these four models are directly taken from GraphReg. **f**) Comparing EPCOT’s within-cell type and cross-cell type prediction with CAGE-seq prediction using the averaged CAGE-seq signals. Here we average ten cell types except the one to be predicted. Within-cell type prediction is performed on four training cell types and cross-cell type prediction is performed on the remaining cell types. Correlation is calculated across genomic bins associated with gene TSS in testing genomic regions, CAGE-seq signals are log-transformed with a pseudo-count of 1. EPCOT achieves higher correlation for majority of the cell types.

**Extended Data Fig.5:**
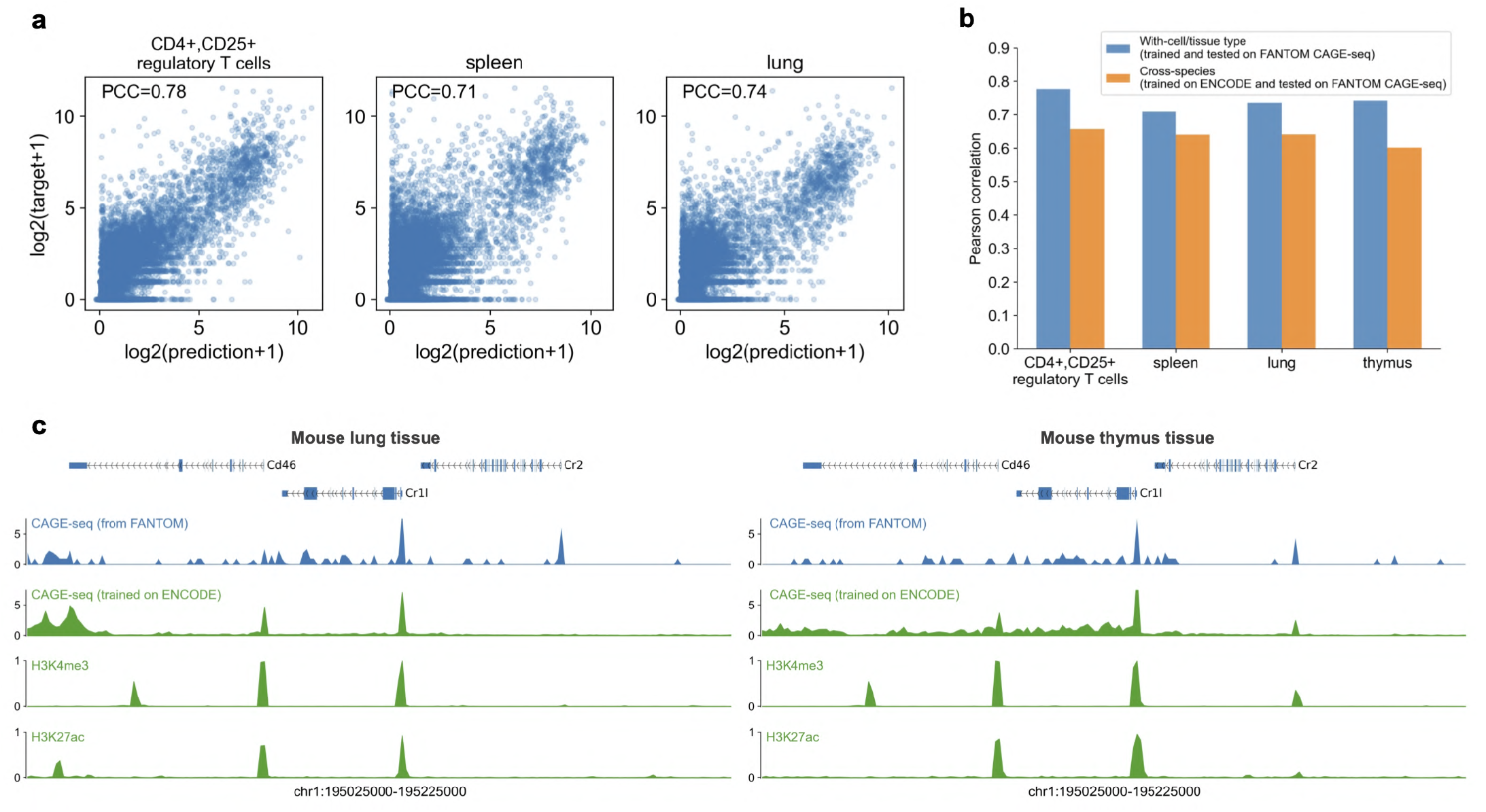
EPCOT predicts cross-species CAGE-seq for mouse cell types and tissues. **a)** Within-cell type CAGE-seq prediction on mouse cell types and tissues. The predicted and target CAGE-seq values on each 1kb genomic bin in the testing regions along with Pearson correlation over all genomic bins, are provided. b) Pearson correlation in with-cell type and cross-species prediction. In cross-species prediction, the model trained on human GM12878, K562, HUVEC, and IMR-90 CAGE-seq from ENCODE is tested on mouse cell types or tissues. The mouse CAGE-seq is collected from FANTOM. **c)** Cross-species CAGE-seq and epigenomic feature prediction for two mouse tissues in the same genomic region with Fig.3h. The epigenomic features are predicted from the human pre-training model.

**Extended Data Fig.6:**
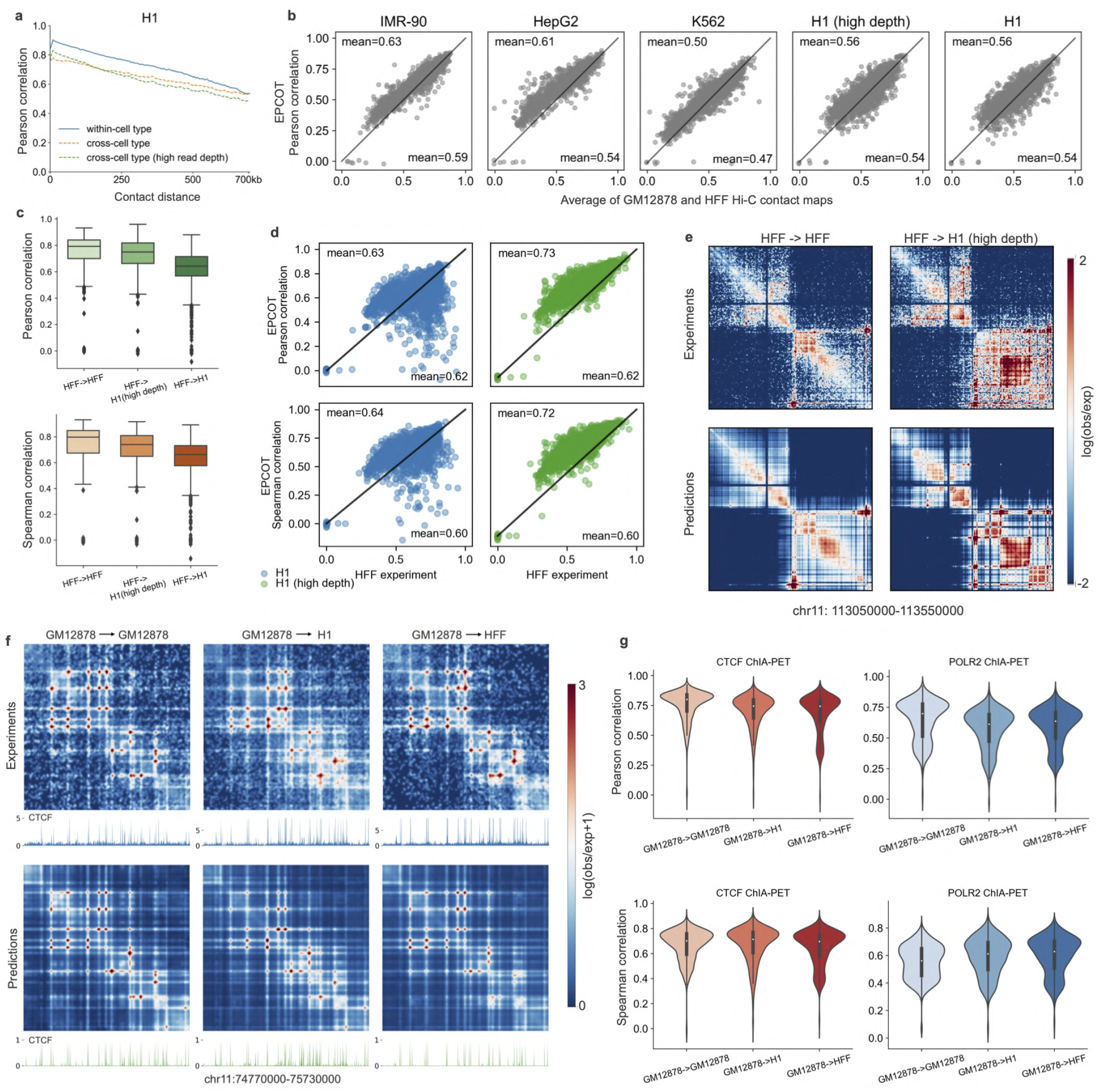
EPCOT predicts cross-cell type chromatin contact maps. **a)** Distance-stratified Pearson correlation of within-cell type and cross-cell type prediction of H1 Hi-C contact maps. In addition to the low read depth H1 DNase-seq, the cross-cell type prediction is also performed on another H1 DNase-seq profile at much higher read depth which is also available from ENCODE. **b)** Comparisons between EPCOT’s cross-cell type and cross-chromosome Hi-C prediction and the prediction by averaging two training cell types’ contact maps. Each dot represents Pearson correlation in a 1Mb testing genomic region. **c)** Within-cell type and cross-cell type prediction performance in 1kb-resolution Micro-C contact map prediction. The model trained on HFF is tested on HFF and another two H1’s DNase-seq at different read depths. **d)** Comparisons between EPCOT’s cross-cell type and cross-chromosome prediction of H1’s Micro-C contact maps and the prediction using HFF’s Micro-C. Each dot represents a correlation score in a 500kb testing region. **e)**An example region to show cross-cell type prediction of 1kb-resolution Micro-C contact map. Micro-C prediction for other cell types and tissues is shown in Supplementary Figure 2. **f)** EPCOT predicts cell-type specific CTCF ChIA-PET. The model trained on GM12878 is used to predict three cell lines. An example region of predicted and target ChIA-PET is given, and the predicted CTCF binding activities and CTCF tracks in each testing cell line are also provided. **g)** Within-cell type and cross-cell type prediction performance for CTCF and POLR2A ChIAPET. The model trained on GM12878 is tested on itself and other two cell lines. Pearson correlation is calculated for each test genomic region.

**Extended Data Fig.7:**
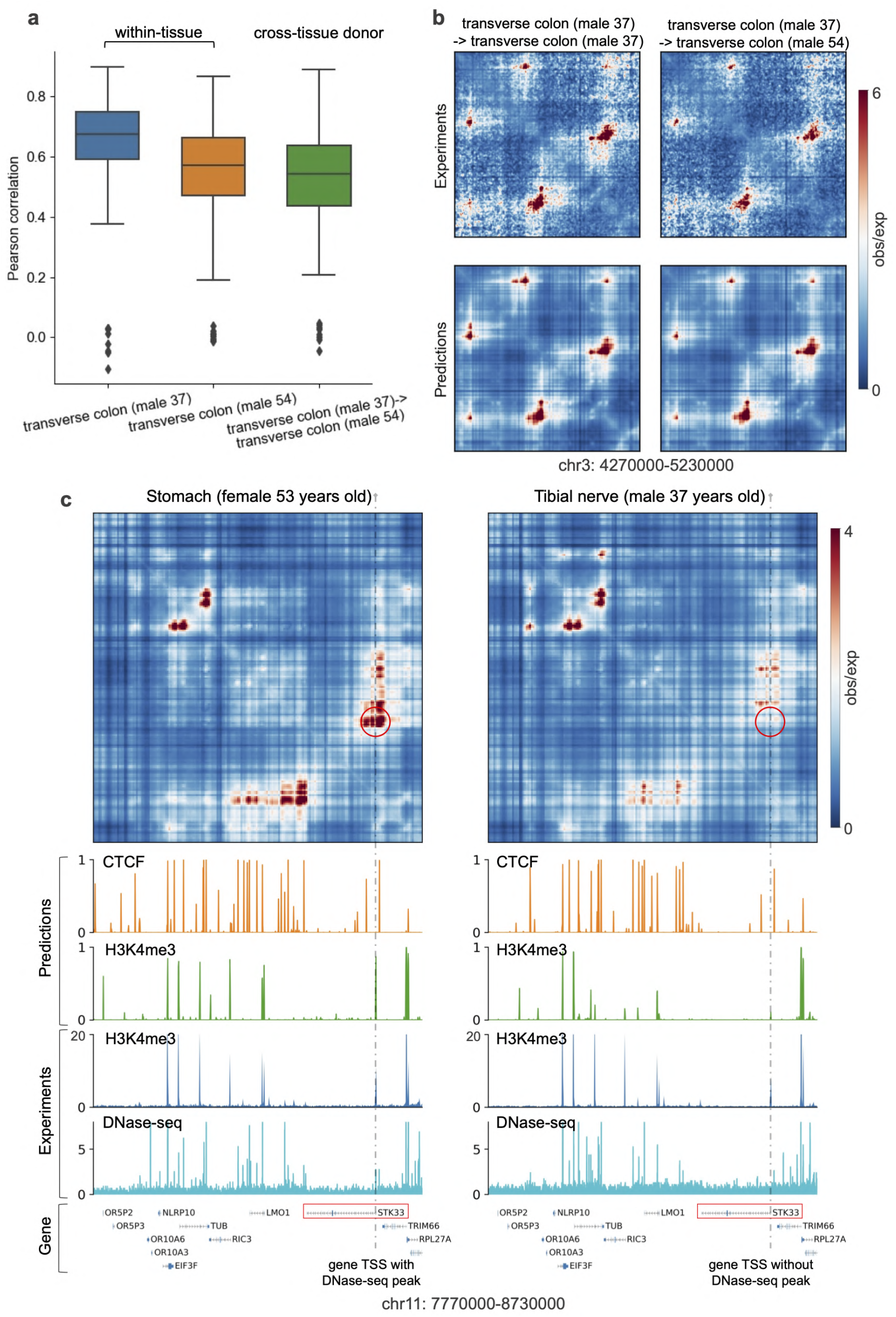
EPCOT predicts chromatin contact maps for GTEx tissues. **a)** Pearson correlation of two tissue donors under within-tissue and cross-tissue donor prediction. **b)** An example region to show within-tissue and cross-tissue donor prediction. **c)** Model trained on traverse colon tissue (male 37 years old) predicts cross-tissue Hi-C contact maps. The difference between predicted chromatin contacts of stomach and tibial nerve tissues (inside red circles) can result from the the difference of DNase-seq at STK33 gene TSS.

**Extended Data Fig.8:**
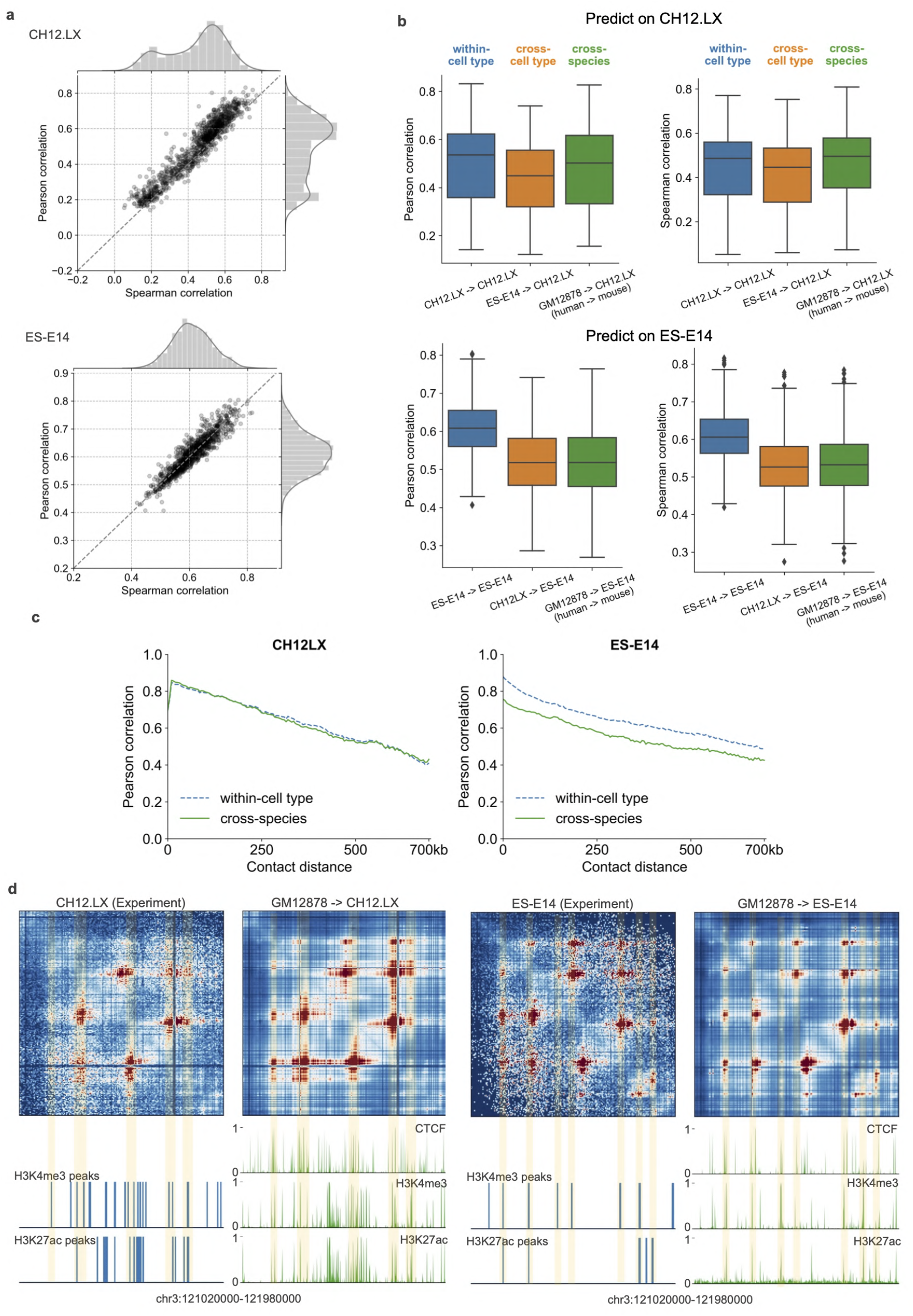
An EPCOT model trained on human cell line GM12878 predicts Hi-C contact maps for two mouse cell lines ES-E14 and CH12.LX under different experiment settings. **a)** Pearson correlation and Spearman correlation in test chromosomes 3 and 11 of two mouse cell lines under within-cell type prediction. **b)** Pearson correlation and Spearman correlation in within-cell type, cross-cell type and cross-species prediction. **c)** Distance-stratified Pearson correlation in within-cell type and cross-species prediction. The model trained on human GM12878 cell line is tested on mouse CH12.LX and ES-E14 cell lines in cross-species prediction. **d)** Cross-species Hi-C, CTCF, H3K4me3, and H3K27ac prediction on two mouse cell lines. The same genomic region in Fig.4e is provided, the mouse Hi-C is predicted by the model trained on human GM12878 cell line, and the epigenomic profiles are predicted by the human pre-training model.

**Extended Data Fig.9:**
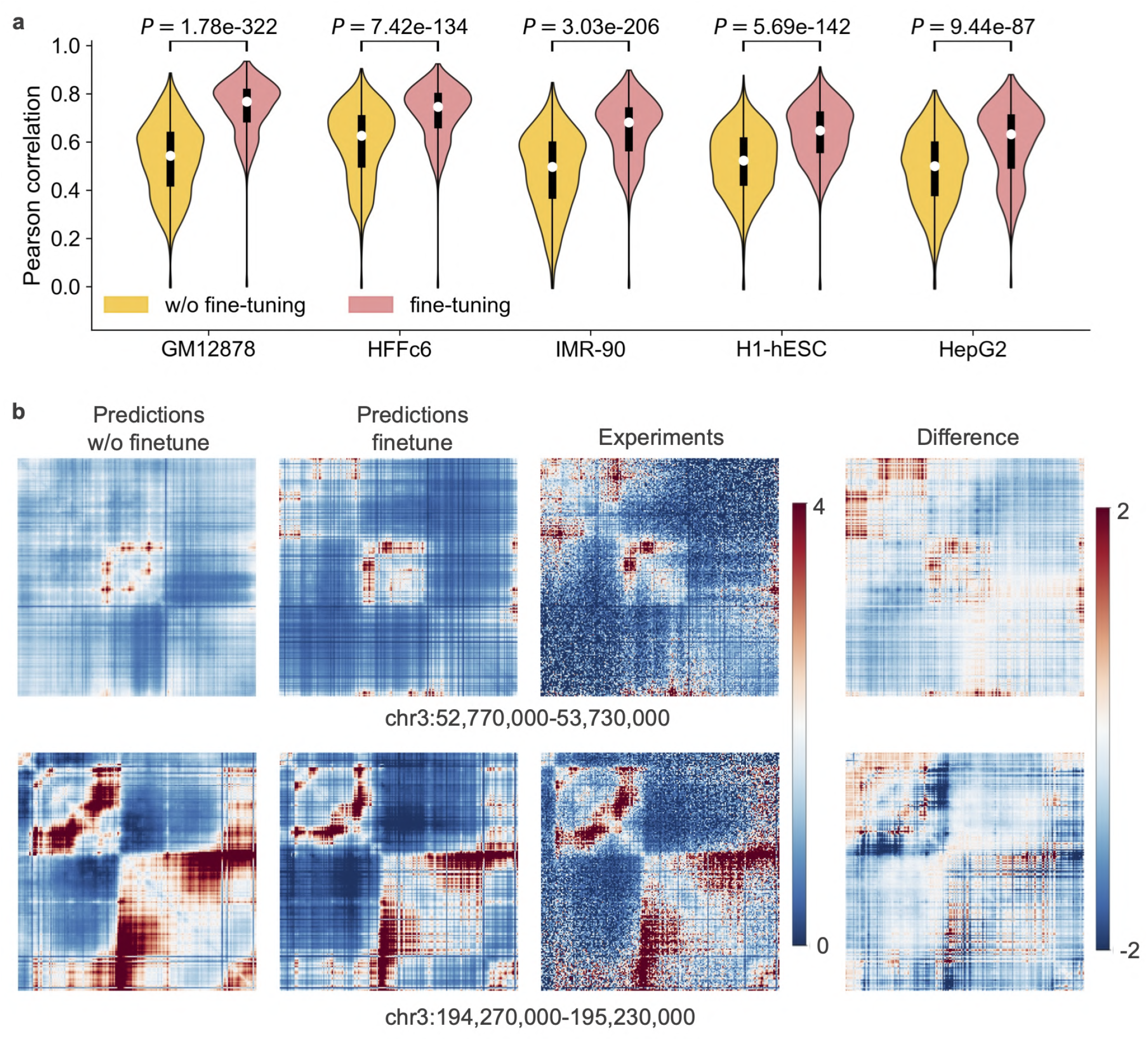
Fine-tuning significantly improves Hi-C contact map prediction. **a)** Comparing Pearson correlation of predicted and target Hi-C in 1Mb genomic regions from the test chromosomes when the pretraining model is fine-tuned or not. **b)** Comparing IMR-90 Hi-C contact maps predicted by EPCOT with or without fine-tuning in two genomic regions. In the top genomic region, the contact matrices predicted without fine-tuning are dissimilar to the target comparing to the prediction with fine-tuning. In the bottom genomic region, EPCOT with fine-tuning and without fine-tuning predicts the target contact map well. EPCOT with fine-tuning predicts salient regions better than EPCOT without fine-tuning by comparing the difference between their predicted contact maps with the target in these two scenarios.

**Extended Data Fig.10:**
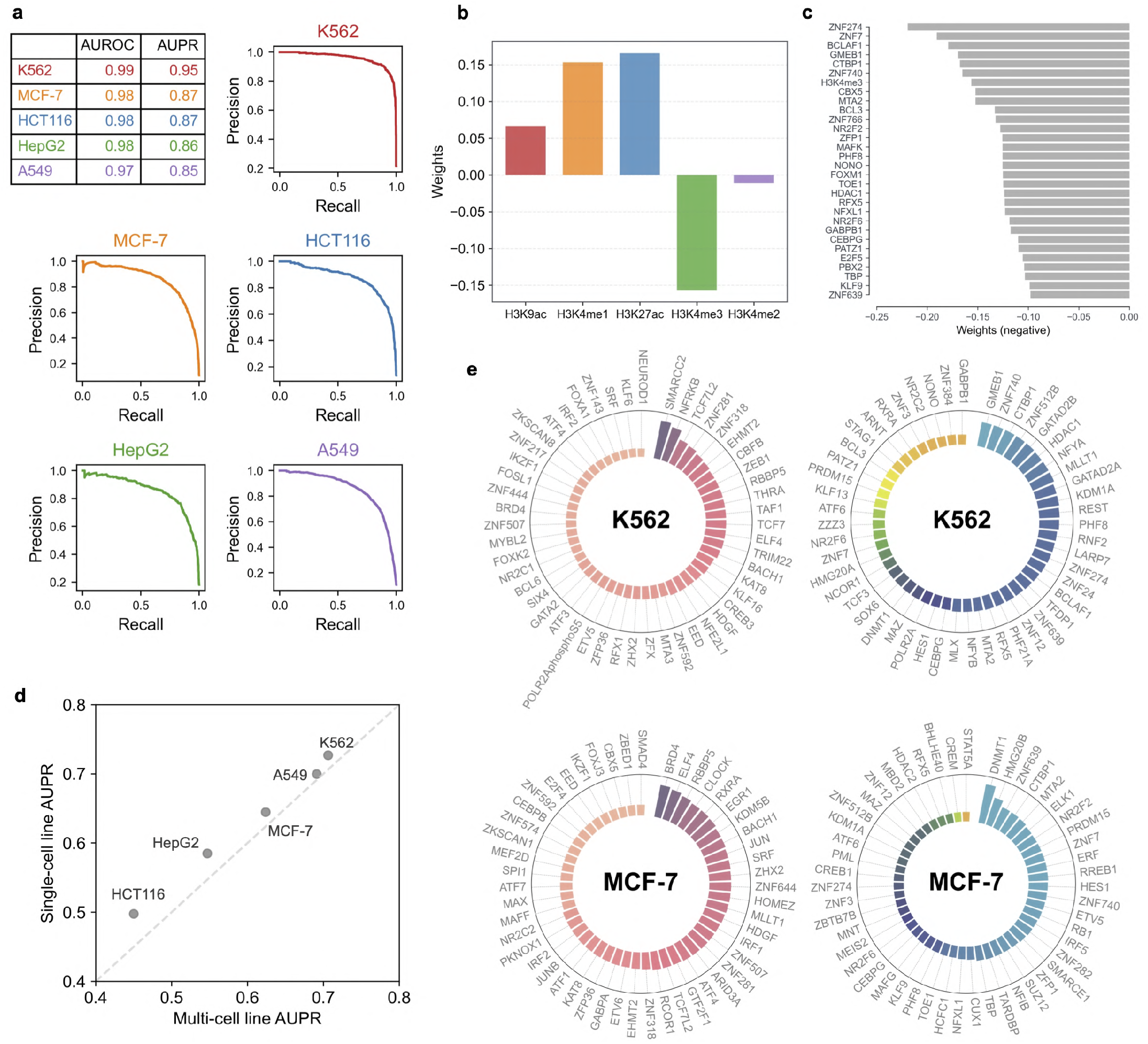
ECPOT accurately predicts active enhancers and characterizes the contributions of epigenomic features. **a**) Enhancer activity prediction performance in a 1-to-10 ratio of positive to negative samples. The AUPR and AUROC scores of five cell lines are shown in a table, and the AUPR curves are also provided. **b**) Weights for five histone marks in the logistic regression (LR) model to predict enhancer activity. **c**) Top 30 negative weights in the LR model to predict enhancer activity in five cell lines. **d**) The LR model trained on individual cell line outperforms the model trained on all five cell lines, which indicates the possibility of cell-type specific relationships between epigenomic features and enhancer activity. **e**) Top 50 positive and negative weights for TFs in the LR model of two cell lines, which characterizes TFs’ cell-type specific relationship to enhancer activity.

**Extended Data Fig.11:**
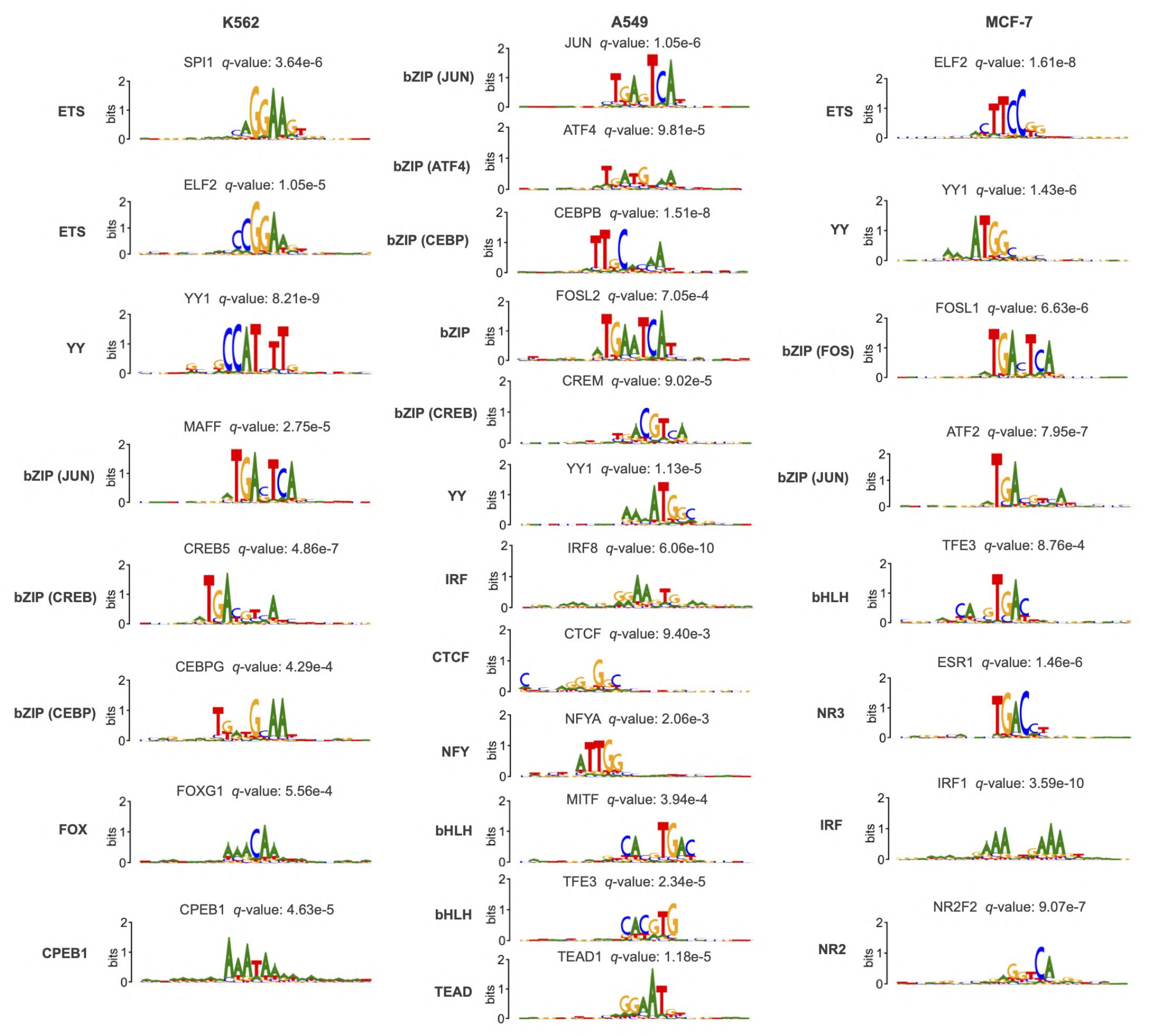
Cell-type specific sequence patterns in active enhancer regions of three cell lines. For each sequence pattern, the most matched TF motifs from HOCOMOCO along with *q*-value (lower indicates better match) using Tomtom are provided.

## Supplementary information

### DNase-seq processing

EPCOT predicts cell-type specific epigenomic features, gene expression, chromatin contact maps, and enhancer activity with the only cell-type specific input of DNase-seq, so we use deepTools [64] to normalize DNase in all used cell types: we download the bam files from ENCODE [40] and merge different replicates, which are then converted into bigWig files and normalized by using deepTools [64] bamCoverage’s RPGC normalization with the parameter binSize 1.

### Epigenomic feature prediction (EFP)

In the EFP task, we first labelled 1kb genomic sequences in a way similar to DeepSEA that if more than 100bp of a 1kb bin was in the peak region of an epigenomic feature, then the epigenomic feature in this bin was labelled 1, and 0 otherwise. In this task, we compared EPCOT with two models that also leveraged multi-task training: FactorNet and scFAN. FactorNet used similar model structure to DanQ. scFAN usec similar model structure to DeepSEA, but scFAN only used half number of kernels in three convolutional layers compared to DeepSEA. To be consistent with EPCOT, we only used the forward strand of DNA sequence and DNase-seq as input. Chromosomes 2, 10, and 21 were used for testing, and the remaining chromosomes were used for validation and training. In the training, we only used genomic sequences which had binding activity for at least one epigenomic feature, while in the testing, we used all the genomic sequences without genome gaps. In scFAN training, we doubled the number of convolutional kernels to be consistent with DeepSEA, which outperformed the original scFAN architecture in our EFP task.

To train the pre-training model that was used for the downstream tasks, we randomly split 10% input samples for validation, and the remaining samples were all used for training. In addition to a pre-training model trained on DNase-seq, we also provided a model trained on ATAC-seq using the same four cell lines with DNase-seq.

### Gene expression prediction (GEP)

In the within-cell type RNA-seq GEP task, we used the same training, validation and testing genes from chromosomes 1-22 and X in GC-MERGE to compare EPCOT with GC-MERGE. In the cross-cell type RNA-seq GEP task, we first collected RNA-seq and DNase-seq in eight cell types which were both available on ENCODE, and trained EPCOT using four cell types: H1, GM12878, A549, and HeLa-S3, and tested the model on the same test gene sets of four different cell types from chromosomes 1-22 and X with AttentiveChrome and DeepChrome.

In the CAGE-seq GEP task, we downloaded human CAGE-seq bam files from ENCODE [40] and mouse CAGE-seq bam files from FANTOM, then merged different replicates and used the scripts ‘bam cov.py’ and ‘basenji data read.py’ provided by Basenji [4] to convert the bam files into bigWig and process the data. Then, we first compared EPCOT with GraphReg in 5kb-resolution CAGE-seq prediction, the same 10-fold cross-validation was utilized to obtain the prediction results on 20 chromosomes. Due to the computational resource limitation, we split one 6Mb region into six 1Mb regions in the training. In 1kb-resolution CAGE-seq prediction, we used a 250kb window to slide over the whole genome with a step size of 25kb to generate input genomic regions, and used the 250kb genomic region to predict CAGE-seq on the centered 200kb, so there was no overlap for the centered 200kb to be predicted on. Then, we collected CAGE-seq in eleven cell types from ENCODE, and trained EPCOT on four cell types, GM12878, K562, HUVEC, and IMR-90.

### Chromatin organization prediction (COP)

For the Hi-C, Micro-C, and ChIA-PET contact maps of cell types, we downloaded their .hic files from 4DN [62] and obtained the observed and expected contact ratios using Juicebox [65]. To predict Micro-C, we simply took a natural logarithm of observed and expected contact ratios, and clipped to (−2,2), and smoothed with a 2D Gaussian filter (sigma=1, width=5). For ChIA-PET, we took a natural logarithm of observed/expected (O/E) contact values with a pseudo-count of 1, and clipped to (0,5), and smoothed with a 2D Gaussian filter (sigma=1, width=5). While in the 5kb-resolution Hi-C contact map prediction of six cell lines (GM12878, K562, IMR-90, H1, HFF, and HepG2), we directly predicted the observed and expected contact ratios without any smoothing. Hi-C of two tissue donors, transverse colon (male 37 years old) and transverse colon (male 54 years old), were downloaded from ENCODE and processed using 4DN pipeline, which were then O/E-normalized and smoothed with a 2D Gaussian filter (sigma=0.6, width=5) due to the low read depth.

In both the 5kb-resolution Hi-C COP and ChIA-PET prediction, we predicted the upper triangle of the contact matrices in the centered 960kb of 1Mb genomic regions. The input genomic regions were generated by using a 1Mb window to slide over the whole genomic sequence with a step size of 250kb, and the regions without genome gaps were kept. Chromosomes 3, 11, and 17 were used for testing, and chromosomes 9 and 16 were used for validation, and the remaining chromosomes were training chromosomes. To compare with Epiphany and DeepC, we generated the 1Mb testing regions with a step size of 50kb and predicted the upper triangle of the contact matrices for each testing region, and simply took an average of the predicted chromatin contacts in overlapping regions. Then we can predict the distance-stratified Pearson correlation and Spearman correlation up to 910kb.

In the Micro-C COP task, to compare with Akita, we predicted our processed Micro-C contact maps in 2kb resolution and used 1Mb genomic regions to predict the contact matrices in the centered 960kb. Here, the same training/validation/testing split was used. In the 1kb-resolution Micro-C COP, we used a 600kb genomic region to predict the upper triangle of contact matrix in the centered 500kb region, and the same training/validation/testing split was used as the split in Hi-C COP task.

### Enhancer activity prediction (EAP)

In the EAP task, we labelled active enhancers in a way that if more than 100bp of a 1kb bin was in the region of candidate enhancers, STARR-seq peaks and H3K27ac peaks, then this bin is labelled as an active enhancer. In this task, chromosomes 2, 10, and 21 were used for model testing, and the remaining chromosomes were used for training and validation. Since the number of negatives was far more than the number of positives (Supplementary Table 4), to train the model, we simply down-sampled the negatives by randomly choosing ten times more negatives than positives.

### Sequence pattern generation

To generate the sequence binding patterns of TFs, we first calculate the attribution scores of DNA sequence in the pre-training model. For each TF, we first randomly selected 180 input genomic regions which are bound by this TF and received corresponding predicted scores greater than 0.4 in the same cell line. Then a gradient-based attribution method ‘*gradient×input*’ was adopted to calculate the attribution scores by multiplying the absolute value of the gradient of predicted score with respect to the genomic sequences with the one-hot encoded sequence. Once obtaining the attribution scores of each TF, its sequence patterns were generated by using TF-MoDISco [30] which first identifies seqlets (high-contribution regions), then splits the seqlets into two metaclusters, and clusters the seqlets in each metacluster, and aggregate seqlet clusters into sequence patterns. For each TF, we simply used same setting to run TF-MoDISco (sliding window size of 20, filter out clusters with less than 45 seqlets) except for ADNP and CHD4 whose generated sequence patterns seem longer than others.

The cell-type specific sequence patterns in active enhancers are generated from attribution scores of 600 active enhancer sequences which receive predicted scores greater than 0.4 in each investigated cell line. Then, TF-MoDISco is run with sliding window size of 16 and filter out clusters with less than 60 seqlets.

**Supplementary Figure 1:**
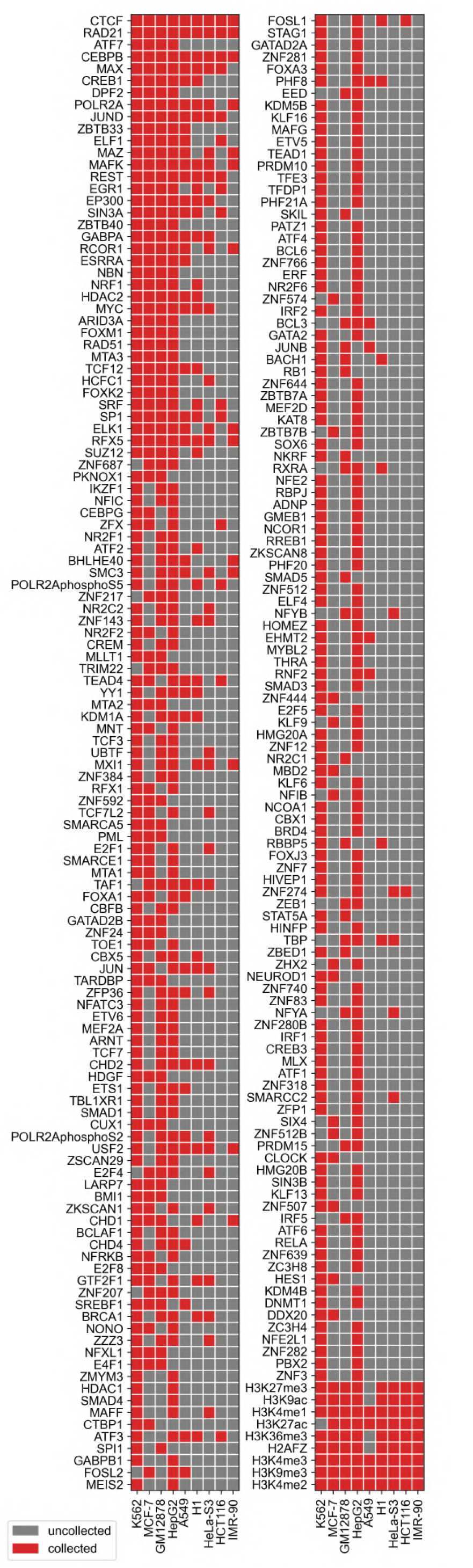
Cell types and epigenomic features used for the pre-training model’s training and testing.

**Supplementary Table 1:**
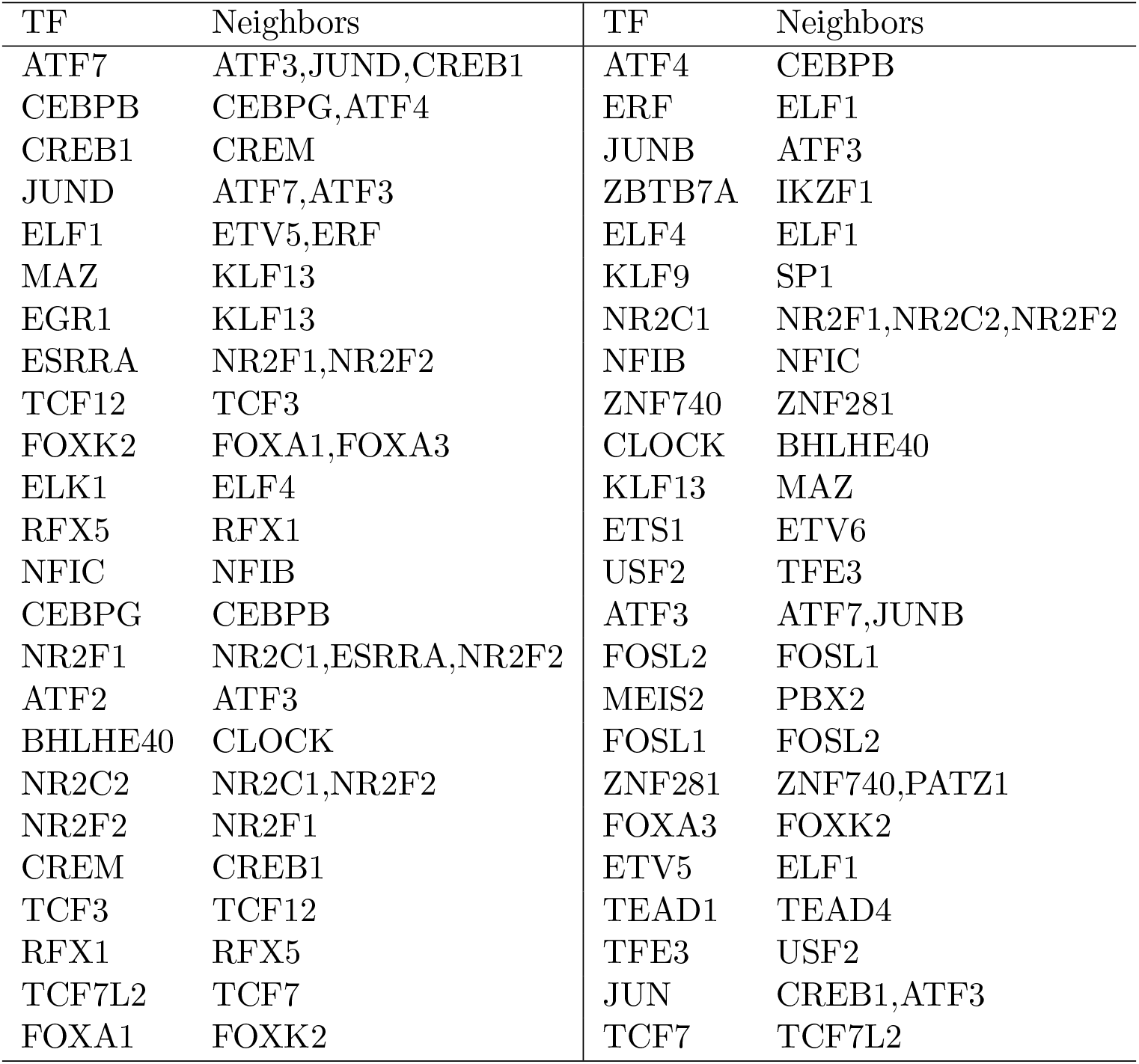
A list of TFs and their embedding neighbors whose binding motifs are in the same cluster [28, 29]

| TF | Neighbors | TF | Neighbors |
| --- | --- | --- | --- |
| ATF7 | ATF3,JUND,CREB1 | ATF4 | CEBPB |
| CEBPB | CEBPG,ATF4 | ERF | ELF1 |
| CREB1 | CREM | JUNB | ATF3 |
| JUND | ATF7,ATF3 | ZBTB7A | IKZF1 |
| ELF1 | ETV5,ERF | ELF4 | ELF1 |
| MAZ | KLF13 | KLF9 | SP1 |
| EGR1 | KLF13 | NR2C1 | NR2F1,NR2C2,NR2F2 |
| ESRRA | NR2F1,NR2F2 | NFIB | NFIC |
| TCF12 | TCF3 | ZNF740 | ZNF281 |
| FOXK2 | FOXA1,FOXA3 | CLOCK | BHLHE40 |
| ELK1 | ELF4 | KLF13 | MAZ |
| RFX5 | RFX1 | ETS1 | ETV6 |
| NFIC | NFIB | USF2 | TFE3 |
| CEBPG | CEBPB | ATF3 | ATF7,JUNB |
| NR2F1 | NR2C1,ESRRA,NR2F2 | FOSL2 | FOSL1 |
| ATF2 | ATF3 | MEIS2 | PBX2 |
| BHLHE40 | CLOCK | FOSL1 | FOSL2 |
| NR2C2 | NR2C1,NR2F2 | ZNF281 | ZNF740,PATZ1 |
| NR2F2 | NR2F1 | FOXA3 | FOXK2 |
| CREM | CREB1 | ETV5 | ELF1 |
| TCF3 | TCF12 | TEAD1 | TEAD4 |
| RFX1 | RFX5 | TFE3 | USF2 |
| TCF7L2 | TCF7 | JUN | CREB1,ATF3 |
| FOXA1 | FOXK2 | TCF7 | TCF7L2 |

**Supplementary Table 2:** 115 pairs of TFs identified by five nearest neighbors using cosine similarity, receive confidence scores greater than 0.4 from STRING database.

| TF interactions | Score | TF interactions | Score | TF interactions | Score |
| --- | --- | --- | --- | --- | --- |
| MTA1 ⟷ RCOR1 | 0.68 | ATF3 ⟷ JUND | 0.974 | ATF7 ⟷ JUND | 0.604 |
| E2F4 ⟷ GATA2 | 0.616 | RB1 ⟷ TFDP1 | 0.998 | CTBP1 ⟷ TCF7 | 0.841 |
| MYC ⟷ ZFP1 | 0.87 | ATF3 ⟷ JUNB | 0.952 | HDAC2 ⟷ RCOR1 | 0.999 |
| SMAD4 ⟷ THRA | 0.546 | GABPB1 ⟷ NRF1 | 0.875 | BRCA1 ⟷ E2F4 | 0.713 |
| GATA2 ⟷ HDAC1 | 0.54 | JUN ⟷ ZHX2 | 0.415 | FOSL1 ⟷ FOSL2 | 0.827 |
| EGR1 ⟷ IRF5 | 0.913 | MAFG ⟷ MAFK | 0.907 | EHMT2 ⟷ PHF21A | 0.472 |
| FOSL1 ⟷ MAFG | 0.44 | CBX1 ⟷ TBL1XR1 | 0.908 | MAFK ⟷ RFX5 | 0.513 |
| ATF7 ⟷ CREB1 | 0.595 | SMAD4 ⟷ TCF3 | 0.919 | NFIB ⟷ NFIC | 0.697 |
| RBBP5 ⟷ SMARCA5 | 0.453 | ZBTB7A ⟷ ZBTB7B | 0.453 | EP300 ⟷ SPI1 | 0.984 |
| TCF12 ⟷ TCF3 | 0.652 | NR2C1 ⟷ NR2C2 | 0.688 | EHMT2 ⟷ MTA3 | 0.666 |
| CBX5 ⟷ TFDP1 | 0.907 | JUN ⟷ MAFK | 0.561 | CHD1 ⟷ SMARCE1 | 0.447 |
| JUNB ⟷ SP1 | 0.93 | HCFC1 ⟷ ZNF143 | 0.49 | MYC ⟷ POLR2A | 0.828 |
| SMARCC2 ⟷ SMARCE1 | 0.999 | IRF1 ⟷ IRF2 | 0.935 | CTCF ⟷ SMC3 | 0.96 |
| CREB1 ⟷ CREM | 0.797 | CREB1 ⟷ JUN | 0.998 | BRCA1 ⟷ NCOR1 | 0.43 |
| CEBPB ⟷ CEBPG | 0.912 | E2F1 ⟷ EED | 0.915 | E2F5 ⟷ FOXM1 | 0.412 |
| HDAC1 ⟷ NCOR1 | 0.999 | HDAC1 ⟷ ZNF217 | 0.663 | JUND ⟷ TEAD4 | 0.453 |
| ATF2 ⟷ ATF3 | 0.723 | KLF6 ⟷ ZFP36 | 0.44 | ATF4 ⟷ CEBPB | 0.976 |
| MYC ⟷ YY1 | 0.983 | RBBP5 ⟷ TCF7L2 | 0.602 | NCOR1 ⟷ NR2F2 | 0.844 |
| KDM5B ⟷ PHF8 | 0.699 | NFYB ⟷ SREBF1 | 0.668 | GATA2 ⟷ HMG20B | 0.41 |
| CREB1 ⟷ MEF2A | 0.418 | CREB1 ⟷ FOSL2 | 0.747 | ATF3 ⟷ ATF7 | 0.411 |
| BRCA1 ⟷ JUN | 0.892 | GATAD2A ⟷ HMG20A | 0.67 | HDAC2 ⟷ STAT5A | 0.414 |
| CHD4 ⟷ KDM1A | 0.959 | CEBPB ⟷ CREB1 | 0.975 | RAD21 ⟷ SMC3 | 0.999 |
| BRD4 ⟷ SMC3 | 0.487 | MAFG ⟷ NFE2L1 | 0.997 | EP300 ⟷ FOSL2 | 0.496 |
| DNMT1 ⟷ SUZ12 | 0.911 | MYC ⟷ TFDP1 | 0.94 | IKZF1 ⟷ MEIS2 | 0.427 |
| BRCA1 ⟷ RELA | 0.96 | BHLHE40 ⟷ CLOCK | 0.958 | MEIS2 ⟷ PBX2 | 0.949 |
| CBX1 ⟷ PHF8 | 0.535 | MTA2 ⟷ MTA3 | 0.947 | ETS1 ⟷ ETV6 | 0.545 |
| DNMT1 ⟷ MTA1 | 0.883 | PBX2 ⟷ PKNOX1 | 0.899 | MEF2A ⟷ TCF3 | 0.685 |
| ATF3 ⟷ FOSL2 | 0.614 | FOSL2 ⟷ SMAD3 | 0.512 | ATF4 ⟷ EP300 | 0.969 |
| CTCF ⟷ RAD21 | 0.996 | GATA2 ⟷ NR2F2 | 0.743 | SMC3 ⟷ STAG1 | 0.999 |
| NCOR1 ⟷ RBPJ | 0.995 | MAFF ⟷ MAFK | 0.91 | CTCF ⟷ STAG1 | 0.863 |
| RBPJ ⟷ SMAD3 | 0.448 | KDM5B ⟷ MYC | 0.976 | EED ⟷ SMARCA5 | 0.451 |
| DPF2 ⟷ ZC3H8 | 0.417 | BCLAF1 ⟷ CHD1 | 0.456 | RBBP5 ⟷ TAF1 | 0.944 |
| MXI1 ⟷ SIN3B | 0.724 | SMAD4 ⟷ TFE3 | 0.955 | CBX5 ⟷ H2AFZ | 0.622 |
| ATF3 ⟷ MAFG | 0.54 | HDAC2 ⟷ SIN3A | 0.999 | NFE2 ⟷ ZNF24 | 0.419 |
| RELA ⟷ SMAD3 | 0.57 | ATF3 ⟷ JUN | 0.999 | ADNP ⟷ CHD4 | 0.896 |
| MYC ⟷ RB1 | 0.866 | ATF3 ⟷ ATF4 | 0.981 | TEAD1 ⟷ TEAD4 | 0.955 |
| RAD21 ⟷ STAG1 | 0.999 | ETV6 ⟷ ZNF384 | 0.563 | EHMT2 ⟷ REST | 0.863 |
| MAFG ⟷ NFE2 | 0.99 | E2F4 ⟷ TAF1 | 0.53 | CREB3 ⟷ NFE2 | 0.402 |
| MAX ⟷ SIN3A | 0.97 | | | | |

**Supplementary Table 3:** Model performance comparisons in 5kb-resolution Hi-C and 2kb-resolution Micro-C contact map prediction. The Pearson correlation scores and Spearman correlation scores of Epiphany, DeepC, and Akita are reported from their papers.

| 5kb-resolution Hi-C contact map prediction |  |  |
| --- | --- | --- |
|  | Pearson correlation | Spearman correlation |
| Epiphany (GM12878 OE-normalized from HiC-DC+) | 0.563* | 0.527* |
| DeepC (GM12878 unsmoothed percentile normalization) | 0.36* | - |
| EPCOT w/ Transformer (GM12878 OE-normalized) | **0.79\*** | **0.79\*** |
| DeepC (K562 unsmoothed percentile normalization) | 0.28* | - |
| EPCOT w/ Transformer (K562 OE-normalized) | **0.64\*** | **0.58\*** |
| EPCOT w/ Transformer (GM12878 OE-normalized) | **0.741**<sup>◇</sup> | **0.711**<sup>◇</sup> |
| EPCOT w/ LSTM (GM12878 OE-normalized) | 0.731<sup>◇</sup> | 0.706<sup>◇</sup> |
| EPCOT w/ Transformer (IMR-90 OE-normalized) | **0.652**<sup>◇</sup> | **0.602**<sup>◇</sup> |
| EPCOT w/ LSTM (IMR-90 OE-normalized) | 0.639<sup>◇</sup> | 0.587<sup>◇</sup> |
| EPCOT w/ Transformer (HFF OE-normalized) | **0.721**<sup>◇</sup> | **0.683**<sup>◇</sup> |
| EPCOT w/ LSTM (HFF OE-normalized) | 0.661<sup>◇</sup> | 0.627<sup>◇</sup> |
| EPCOT w/ Transformer (H1-hESC OE-normalized) | **0.636**<sup>◇</sup> | **0.604**<sup>◇</sup> |
| EPCOT w/ LSTM (H1-hESC OE-normalized) | 0.536<sup>◇</sup> | 0.513<sup>◇</sup> |
| EPCOT w/ Transformer (HepG2 OE-normalized) | **0.596**<sup>◇</sup> | **0.535**<sup>◇</sup> |
| EPCOT w/ LSTM (HepG2 OE-normalized) | 0.586<sup>◇</sup> | 0.529<sup>◇</sup> |
| EPCOT w/ Transformer (K562 OE-normalized) | **0.543**<sup>◇</sup> | **0.494**<sup>◇</sup> |
| Micro-C contact map prediction |  |  |
|  | Pearson correlation | Spearman correlation |
| Akita (H1-hESC in 2048bp resolution) | - | 0.61 |
| Akita (HFF in 2048bp resolution) | - | 0.57 |
| EPCOT w/ Transformer (H1-hESC in 2kb resolution) | **0.82** | **0.79** |
| EPCOT w/ Transformer (HFF in 2kb resolution) | **0.78** | **0.81** |

\* Average, distance-stratified Pearson correlation/Spearman correlation.
<sup>◇</sup> Average Pearson correlation/Spearman correlation for each 1Mb genomic region in the test chromosomes

**Supplementary Figure 2:**
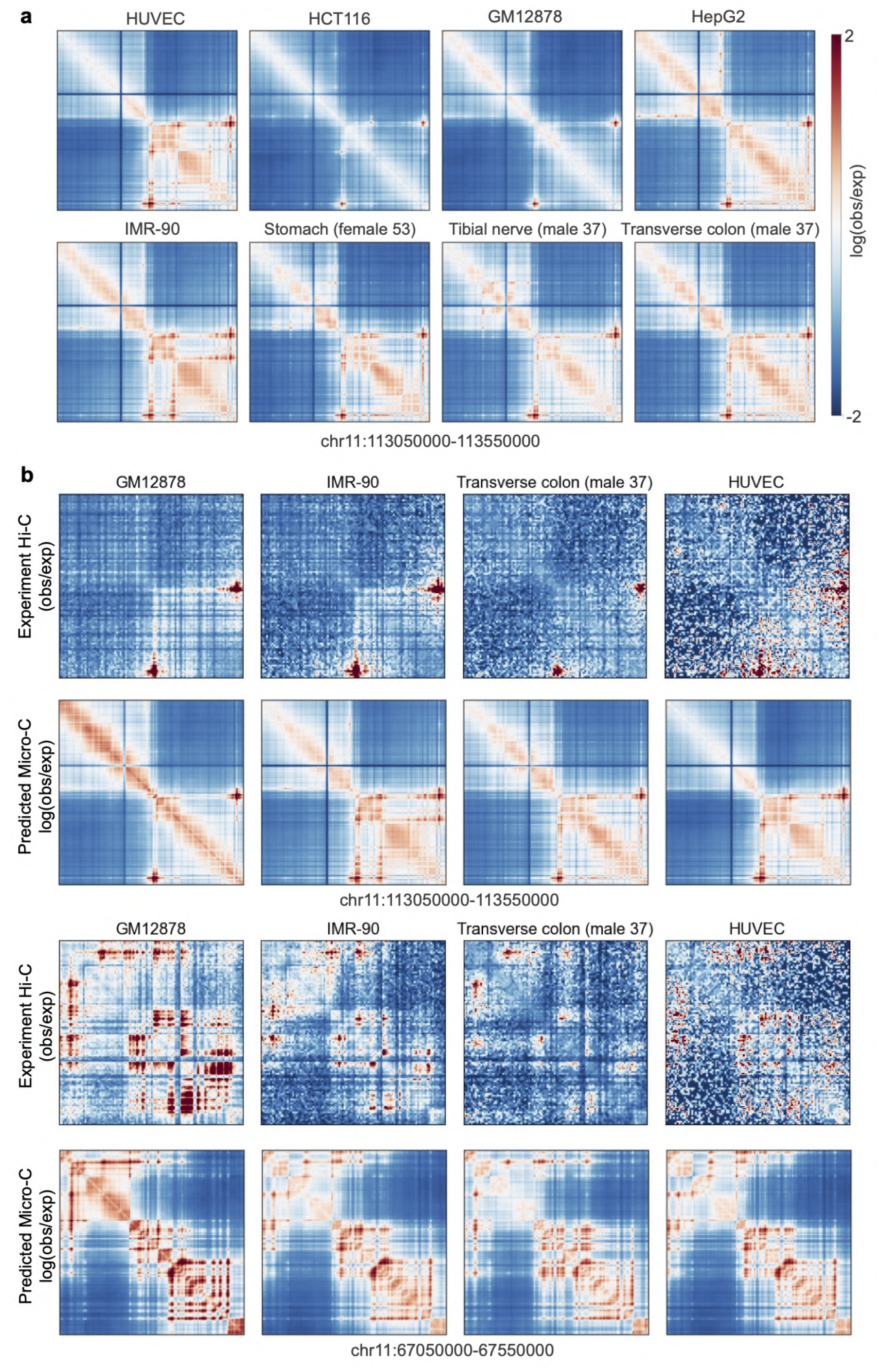
EPCOT predicts cross-cell and tissue type Micro-C contact maps. a) The model trained on HFF cell line is transferred to predict for five cell types and three GTEx tissue donors whose experiment Micro-C contact maps are unavailable. The predicted contact maps in an example region are provided. b) Comparing predicted Micro-C contact maps with available experiment Hi-C contact maps.

**Supplementary Table 4:**
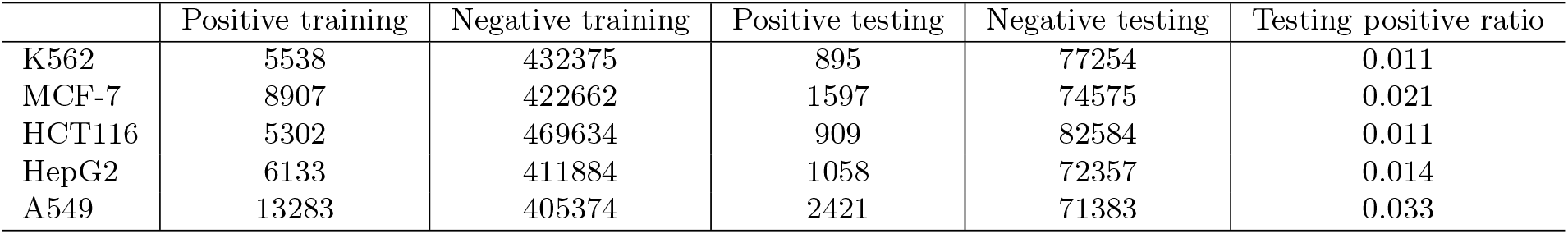
Number of training and testing samples in the positive and negative sets. The testing positive ratio indicates a baseline AUPR score.

|  | Positive training | Negative training | Positive testing | Negative testing | Testing positive ratio |
| --- | --- | --- | --- | --- | --- |
| K562 | 5538 | 432375 | 895 | 77254 | 0.011 |
| MCF-7 | 8907 | 422662 | 1597 | 74575 | 0.021 |
| HCT116 | 5302 | 469634 | 909 | 82584 | 0.011 |
| HepG2 | 6133 | 411884 | 1058 | 72357 | 0.014 |
| A549 | 13283 | 405374 | 2421 | 71383 | 0.033 |

**Supplementary Figure 3:**
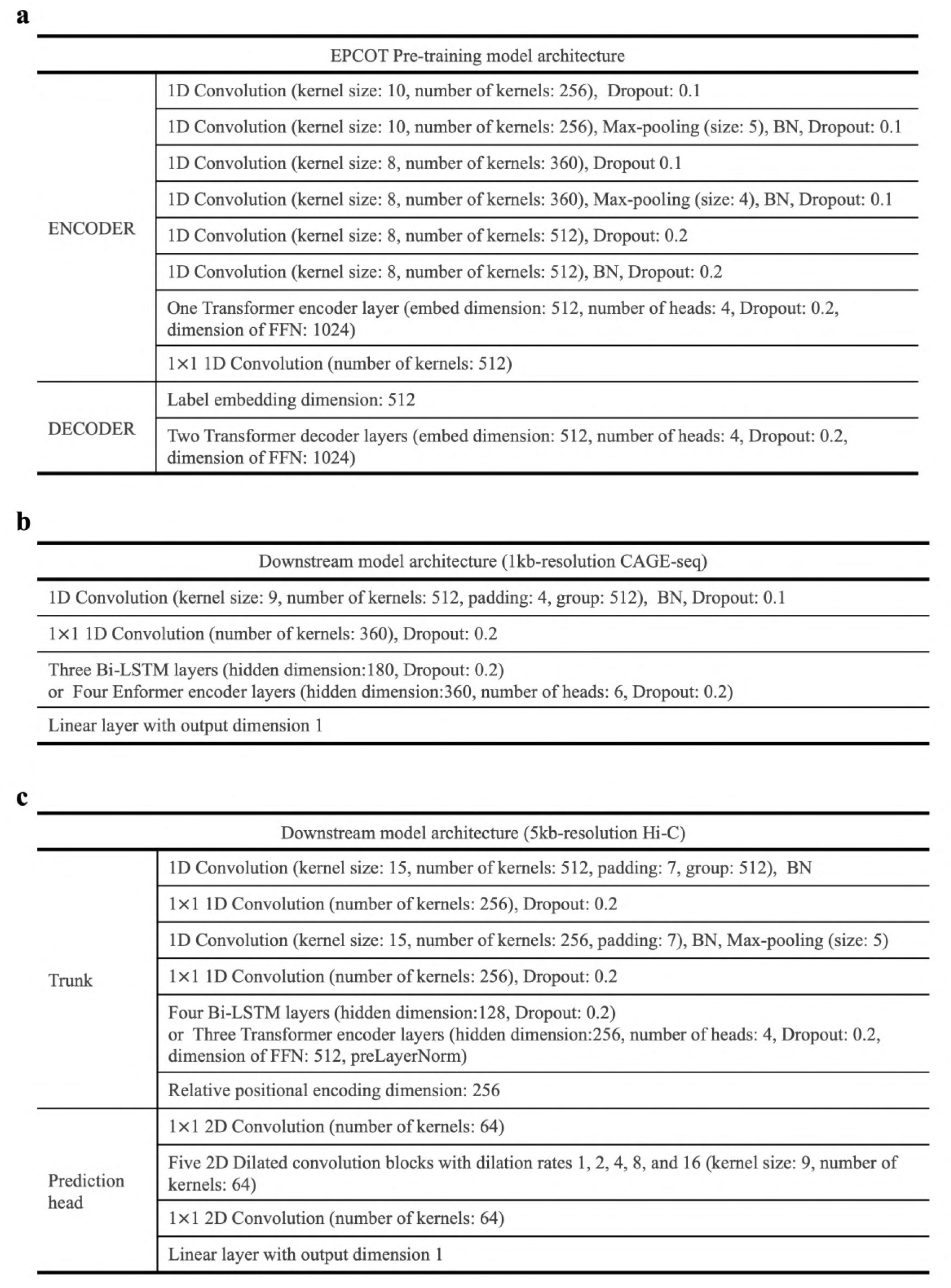
Architectures of EPCOT’s pre-training model and downstream models.

**Supplementary Table 5:** Data sources

| DNase-seq |  |  |  |
| --- | --- | --- | --- |
| HepG2 | ENCSR149XIL | HeLa-S3 | ENCSR959ZXU |
| K562 | ENCSR000EOT | MCF-7 | ENCSR000EPH |
| A549 | ENCSR000ELW | GM12878 | ENCSR000EMT |
| HCT116 | ENCSR000ENM | IMR-90 | ENCSR477RTP |
| HFF | ENCSR672EWY | HUVEC | ENCSR000EOQ |
| H1 | ENCSR000EMU | H1 (high read depth) | ENCSR951BNY |
| skeletal muscle myoblast | ENCSR000EOO | B cell | ENCSR000EMJ |
| transverse colon (male 54) | ENCSR340MRJ | transverse colon (male 37) | ENCSR923JYH |
| stomach (female 53) | ENCSR006IMH | tibial nerve (male 37) | ENCSR484UAU |
| thymus (mouse) | ENCSR000COB | spleen (mouse) | ENCSR000CNY |
| lung (mouse) | ENCSR000CNM | CD4+,CD25+ regulatory T cell | ENCSR000COC |
| CH12.LX (mouse) | ENCSR000CMQ | ES-E14 (mouse) | ENCSR000CMW |
| ATAC-seq |  |  |  |
| GM12878 | ENCSR095QNB | K562 | ENCSR483RKN |
| HepG2 | ENCSR042AWH | MCF-7 | ENCSR422SUG |
| Human CAGE-seq from ENCODE |  |  |  |
| K562 | ENCSR000CJN | GM12878 | ENCSR000CKA |
| HeLa-S3 | ENCSR000CJJ | HepG2 | ENCSR000CJM |
| A549 | ENCSR000CJV | IMR-90 | ENCSR000CKT |
| MCF-7 | ENCSR000CJO | HUVEC | ENCSR000CJE |
| B cell | ENCSR000CKD | CD14+ monocyte | ENCSR000CKW |
| H1 | ENCSR000CJB |  |  |
| Mouse CAGE-seq from FANTOM |  |  |  |
| lung | CNhs10474 | CD4+,CD25+ regulatory T cell | CNhs13913 |
| thymus | CNhs10471 | spleen | CNhs10465 |
| Hi-C |  |  |  |
| H1-hESC | 4DNES2M5JIGV | HFF | 4DNES2R6PUEK |
| IMR-90 | 4DNES1ZEJNRU | K562 | 4DNESI7DEJTM |
| GM12878 | 4DNES3JX38V5 | HepG2 | 4DNESC2DEQIJ |
| ES-E14 | 4DNESU4BQU4G | CH12.LX | 4DNESK95HVFB |
| transverse colon (male 54) | ENCBS682OLO | transverse colon (male 37) | ENCSR295BDK |
| Micro-C |  |  |  |
| H1-hESC | 4DNES21D8SP8 | HFF | 4DNESWST3UBH |
| STARR-seq |  |  |  |
| MCF-7 | ENCSR547SBZ | HCT116 | ENCSR064KUD |
| A549 | ENCSR895FDL | HepG2 | ENCSR135NXN |
| K562 | ENCSR858MPS |  |  |

## References

[1] Jian Zhou and Olga G Troyanskaya. Predicting effects of noncoding variants with deep learning–based sequence model. Nature Methods, 12(10):931–934, 2015.

[2] Daniel Quang and Xiaohui Xie. Factornet: a deep learning framework for predicting cell type specific transcription factor binding from nucleotide-resolution sequential data. Methods, 166:40–47, 2019.

[3] Žiga Avsec, Vikram Agarwal, Daniel Visentin, Joseph R Ledsam, Agnieszka Grabska-Barwinska, Kyle R Taylor, Yannis Assael, John Jumper, Pushmeet Kohli, and David R Kelley. Effective gene expression prediction from sequence by integrating long-range interactions. Nature Methods, 18(10):1196–1203, 2021.

[4] David R Kelley, Yakir A Reshef, Maxwell Bileschi, David Belanger, Cory Y McLean, and Jasper Snoek. Sequential regulatory activity prediction across chromosomes with convolutional neural networks. Genome Research, 28(5):739–750, 2018.

[5] Ritambhara Singh, Jack Lanchantin, Arshdeep Sekhon, and Yanjun Qi. Attend and predict: Understanding gene regulation by selective attention on chromatin. Advances in Neural Information Processing Systems, 30, 2017.

[6] Ritambhara Singh, Jack Lanchantin, Gabriel Robins, and Yanjun Qi. Deepchrome: deep-learning for predicting gene expression from histone modifications. Bioinformatics, 32(17):i639–i648, 2016.

[7] Alireza Karbalayghareh, Merve Sahin, and Christina S Leslie. Chromatin interaction aware gene regulatory modeling with graph attention networks. Genome Research, pages gr–275870, 2022.

[8] Geoff Fudenberg, David R Kelley, and Katherine S Pollard. Predicting 3d genome folding from dna sequence with akita. Nature Methods, 17(11):1111–1117, 2020.

[9] Ron Schwessinger, Matthew Gosden, Damien Downes, Richard C Brown, A Marieke Oudelaar, Jelena Telenius, Yee Whye Teh, Gerton Lunter, and Jim R Hughes. Deepc: predicting 3d genome folding using megabase-scale transfer learning. Nature Methods, 17(11):1118–1124, 2020.

[10] Fan Feng, Yuan Yao, Xue Qing David Wang, Xiaotian Zhang, and Jie Liu. Connecting high-resolution 3d chromatin organization with epigenomics. Nature Communications, 13(1):1–10, 2022.

[11] Jian Zhou. Sequence-based modeling of three-dimensional genome architecture from kilobase to chromosome scale. Nature Genetics, 2022.

[12] Anurag Sethi, Mengting Gu, Emrah Gumusgoz, Landon Chan, Koon-Kiu Yan, Joel Rozowsky, Iros Barozzi, Veena Afzal, Jennifer A Akiyama, Ingrid Plajzer-Frick, et al. Supervised enhancer prediction with epigenetic pattern recognition and targeted validation. Nature Methods, 17(8):807–814, 2020.

[13] Seong Gon Kim, Mrudul Harwani, Ananth Grama, and Somali Chaterji. Ep-dnn: a deep neural networkbased global enhancer prediction algorithm. Scientific Reports, 6(1):1–13, 2016.

[14] Bernardo P de Almeida, Franziska Reiter, Michaela Pagani, and Alexander Stark. Deepstarr predicts enhancer activity from dna sequence and enables the de novo design of enhancers. Nature Genetics, 2022.

[15] Yanrong Ji, Zhihan Zhou, Han Liu, and Ramana V Davuluri. Dnabert: pre-trained bidirectional encoder representations from transformers model for dna-language in genome. Bioinformatics, 37(15):2112–2120, 2021.

[16] ENCODE Project Consortium et al. An integrated encyclopedia of dna elements in the human genome. Nature, 489(7414):57, 2012.

[17] Sandy L Klemm, Zohar Shipony, and William J Greenleaf. Chromatin accessibility and the regulatory epigenome. Nature Reviews Genetics, 20(4):207–220, 2019.

[18] Viraat Y Goel, Miles K Huseyin, and Anders S Hansen. Region capture micro-c reveals coalescence of enhancers and promoters into nested microcompartments. bioRxiv, 2022.

[19] Nicolas Carion, Francisco Massa, Gabriel Synnaeve, Nicolas Usunier, Alexander Kirillov, and Sergey Zagoruyko. End-to-end object detection with transformers. In European Conference on Computer Vision, pages 213–229. Springer, 2020.

[20] Shilong Liu, Lei Zhang, Xiao Yang, Hang Su, and Jun Zhu. Query2label: A simple transformer way to multi-label classification. arXiv preprint arXiv:2107.10834, 2021.

[21] Hongyang Li, Daniel Quang, and Yuanfang Guan. Anchor: trans-cell type prediction of transcription factor binding sites. Genome Research, 29(2):281–292, 2019.

[22] Hongyang Li and Yuanfang Guan. Fast decoding cell type–specific transcription factor binding landscape at single-nucleotide resolution. Genome Research, 31(4):721–731, 2021.

[23] Tareian Cazares, Faiz W Rizvi, Balaji Iyer, Xiaoting Chen, Michael Kotliar, Joseph A Wayman, Anthony Bejjani, Omer Donmez, Benjamin Wronowski, Sreeja Parameswaran, et al. maxatac: genome-scale transcription-factor binding prediction from atac-seq with deep neural networks. bioRxiv, 2022.

[24] Laiyi Fu, Lihua Zhang, Emmanuel Dollinger, Qinke Peng, Qing Nie, and Xiaohui Xie. Predicting transcription factor binding in single cells through deep learning. Science Advances, 6(51):eaba9031, 2020.

[25] Jacob Schreiber, Ritambhara Singh, Jeffrey Bilmes, and William Stafford Noble. A pitfall for machine learning methods aiming to predict across cell types. Genome Biology, 21(1):1–6, 2020.

[26] Laurens Van der Maaten and Geoffrey Hinton. Visualizing data using t-sne. Journal of Machine Learning Research, 9(11), 2008.

[27] Damian Szklarczyk, Annika L Gable, David Lyon, Alexander Junge, Stefan Wyder, Jaime Huerta-Cepas, Milan Simonovic, Nadezhda T Doncheva, John H Morris, Peer Bork, et al. String v11: protein–protein association networks with increased coverage, supporting functional discovery in genome-wide experimental datasets. Nucleic Acids Research, 47(D1):D607–D613, 2019.

[28] Jaime Abraham Castro-Mondragon, Sébastien Jaeger, Denis Thieffry, Morgane Thomas-Chollier, and Jacques Van Helden. Rsat matrix-clustering: dynamic exploration and redundancy reduction of transcription factor binding motif collections. Nucleic Acids Research, 45(13):e119–e119, 2017.

[29] Aziz Khan, Oriol Fornes, Arnaud Stigliani, Marius Gheorghe, Jaime A Castro-Mondragon, Robin Van Der Lee, Adrien Bessy, Jeanne Cheneby, Shubhada R Kulkarni, Ge Tan, et al. Jaspar 2018: update of the open-access database of transcription factor binding profiles and its web framework. Nucleic Acids Research, 46(D1):D260–D266, 2018.

[30] Avanti Shrikumar, Katherine Tian, Žiga Avsec, Anna Shcherbina, Abhimanyu Banerjee, Mahfuza Sharmin, Surag Nair, and Anshul Kundaje. Technical note on transcription factor motif discovery from importance scores (tf-modisco) version 0.5. 6.5. arXiv preprint arXiv:1811.00416, 2018.

[31] Zhenhao Zhang, Fan Feng, and Jie Liu. Characterizing collaborative transcription regulation with a graph-based deep learning approach. PLOS Computational Biology, 18(6):e1010162, 2022.

[32] Žiga Avsec, Melanie Weilert, Avanti Shrikumar, Sabrina Krueger, Amr Alexandari, Khyati Dalal, Robin Fropf, Charles McAnany, Julien Gagneur, Anshul Kundaje, et al. Base-resolution models of transcription-factor binding reveal soft motif syntax. Nature Genetics, 53(3):354–366, 2021.

[33] Veronika Ostapcuk, Fabio Mohn, Sarah H Carl, Anja Basters, Daniel Hess, Vytautas Iesmantavicius, Lisa Lampersberger, Matyas Flemr, Aparna Pandey, Nicolas H Thomä, et al. Activity-dependent neuroprotective protein recruits hp1 and chd4 to control lineage-specifying genes. Nature, 557(7707):739–743, 2018.

[34] Timothy L Bailey, James Johnson, Charles E Grant, and William S Noble. The meme suite. Nucleic Acids Research, 43(W1):W39–W49, 2015.

[35] Jeremy Bigness, Xavier Loinaz, Shalin Patel, Erica Larschan, and Ritambhara Singh. Integrating longrange regulatory interactions to predict gene expression using graph convolutional networks. Journal of Computational Biology, 2022.

[36] Keith Orford, Peter Kharchenko, Weil Lai, Maria Carlota Dao, David J Worhunsky, Adam Ferro, Viktor Janzen, Peter J Park, and David T Scadden. Differential h3k4 methylation identifies developmentally poised hematopoietic genes. Developmental Cell, 14(5):798–809, 2008.

[37] David R Kelley. Cross-species regulatory sequence activity prediction. PLoS Computational Biology, 16(7):e1008050, 2020.

[38] Rui Yang, Arnav Das, Vianne R Gao, Alireza Karbalayghareh, William S Noble, Jeffrey A Bilmes, and Christina S Leslie. Epiphany: predicting hi-c contact maps from 1d epigenomic signals. bioRxiv, 2021.

[39] Shilu Zhang, Deborah Chasman, Sara Knaack, and Sushmita Roy. In silico prediction of high-resolution hi-c interaction matrices. Nature Communications, 10(1):1–18, 2019.

[40] Carrie A Davis, Benjamin C Hitz, Cricket A Sloan, Esther T Chan, Jean M Davidson, Idan Gabdank, Jason A Hilton, Kriti Jain, Ulugbek K Baymuradov, Aditi K Narayanan, et al. The encyclopedia of dna elements (encode): data portal update. Nucleic Acids Research, 46(D1):D794–D801, 2018.

[41] Cosmas D Arnold, Daniel Gerlach, Christoph Stelzer, Lukasz M Boryń, Martina Rath, and Alexander Stark. Genome-wide quantitative enhancer activity maps identified by starr-seq. Science, 339(6123):1074– 1077, 2013.

[42] Tianshun Gao and Jiang Qian. Enhanceratlas 2.0: an updated resource with enhancer annotation in 586 tissue/cell types across nine species. Nucleic Acids Research, 48(D1):D58–D64, 2020.

[43] Biswajyoti Sahu, Tuomo Hartonen, Päivi Pihlajamaa, Bei Wei, Kashyap Dave, Fangjie Zhu, Eevi Kaasinen, Katja Lidschreiber, Michael Lidschreiber, Carsten O Daub, et al. Sequence determinants of human gene regulatory elements. Nature Genetics, 54(3):283–294, 2022.

[44] Nergiz Dogan, Weisheng Wu, Christapher S Morrissey, Kuan-Bei Chen, Aaron Stonestrom, Maria Long, Cheryl A Keller, Yong Cheng, Deepti Jain, Axel Visel, et al. Occupancy by key transcription factors is a more accurate predictor of enhancer activity than histone modifications or chromatin accessibility. Epigenetics & Chromatin, 8(1):1–21, 2015.

[45] Ji-Eun Lee, Young-Kwon Park, Sarah Park, Younghoon Jang, Nicholas Waring, Anup Dey, Keiko Ozato, Binbin Lai, Weiqun Peng, and Kai Ge. Brd4 binds to active enhancers to control cell identity gene induction in adipogenesis and myogenesis. Nature Communications, 8(1):1–12, 2017.

[46] Beibei Liu, Xinhua Liu, Lulu Han, Xing Chen, Xiaodi Wu, Jiajing Wu, Dong Yan, Yue Wang, Shumeng Liu, Lin Shan, et al. Brd4-directed super-enhancer organization of transcription repression programs links to chemotherapeutic efficacy in breast cancer. Proceedings of the National Academy of Sciences, 119(6):e2109133119, 2022.

[47] Gabriel E Zentner, Paul J Tesar, and Peter C Scacheri. Epigenetic signatures distinguish multiple classes of enhancers with distinct cellular functions. Genome Research, 21(8):1273–1283, 2011.

[48] Idit Hazan, Jonathan Monin, Britta AM Bouwman, Nicola Crosetto, and Rami I Aqeilan. Activation of oncogenic super-enhancers is coupled with dna repair by rad51. Cell Reports, 29(3):560–572, 2019.

[49] Suhas SP Rao, Miriam H Huntley, Neva C Durand, Elena K Stamenova, Ivan D Bochkov, James T Robinson, Adrian L Sanborn, Ido Machol, Arina D Omer, Eric S Lander, et al. A 3d map of the human genome at kilobase resolution reveals principles of chromatin looping. Cell, 159(7):1665–1680, 2014.

[50] Jacob Schreiber, Timothy Durham, Jeffrey Bilmes, and William Stafford Noble. Avocado: a multi-scale deep tensor factorization method learns a latent representation of the human epigenome. Genome Biology, 21(1):1–18, 2020.

[51] Jacob Schreiber, Jeffrey Bilmes, and William Stafford Noble. Completing the encode3 compendium yields accurate imputations across a variety of assays and human biosamples. Genome Biology, 21(1):1–13, 2020.

[52] Jason Ernst and Manolis Kellis. Large-scale imputation of epigenomic datasets for systematic annotation of diverse human tissues. Nature Biotechnology, 33(4):364–376, 2015.

[53] Vikram Agarwal and Jay Shendure. Predicting mrna abundance directly from genomic sequence using deep convolutional neural networks. Cell Reports, 31(7):107663, 2020.

[54] Fahad Ullah and Asa Ben-Hur. A self-attention model for inferring cooperativity between regulatory features. Nucleic Acids Research, 49(13):e77–e77, 2021.

[55] Samuel A Lambert, Arttu Jolma, Laura F Campitelli, Pratyush K Das, Yimeng Yin, Mihai Albu, Xiaoting Chen, Jussi Taipale, Timothy R Hughes, and Matthew T Weirauch. The human transcription factors. Cell, 172(4):650–665, 2018.

[56] Tal Ridnik, Gilad Sharir, Avi Ben-Cohen, Emanuel Ben-Baruch, and Asaf Noy. Ml-decoder: Scalable and versatile classification head. arXiv preprint arXiv:2111.12933, 2021.

[57] Ashish Vaswani, Noam Shazeer, Niki Parmar, Jakob Uszkoreit, Llion Jones, Aidan N Gomez, Lukasz Kaiser, and Illia Polosukhin. Attention is all you need. Advances in Neural Information Processing Systems, 30, 2017.

[58] Ilya Loshchilov and Frank Hutter. Decoupled weight decay regularization. arXiv preprint arXiv:1711.05101, 2017.

[59] Sergey Ioffe and Christian Szegedy. Batch normalization: Accelerating deep network training by reducing internal covariate shift. In International Conference on Machine Learning, pages 448–456. PMLR, 2015.

[60] Marina Lizio, Jayson Harshbarger, Hisashi Shimoji, Jessica Severin, Takeya Kasukawa, Serkan Sahin, Imad Abugessaisa, Shiro Fukuda, Fumi Hori, Sachi Ishikawa-Kato, et al. Gateways to the fantom5 promoter level mammalian expression atlas. Genome Biology, 16(1):1–14, 2015.

[61] Anshul Kundaje, Wouter Meuleman, Jason Ernst, Misha Bilenky, Angela Yen, Alireza Heravi-Moussavi, Pouya Kheradpour, Zhizhuo Zhang, Jianrong Wang, Michael J Ziller, et al. Integrative analysis of 111 reference human epigenomes. Nature, 518(7539):317–330, 2015.

[62] Job Dekker, Andrew S Belmont, Mitchell Guttman, Victor O Leshyk, John T Lis, Stavros Lomvardas, Leonid A Mirny, Clodagh C O’shea, Peter J Park, Bing Ren, et al. The 4d nucleome project. Nature, 549(7671):219–226, 2017.

[63] Žiga Avsec, Roman Kreuzhuber, Johnny Israeli, Nancy Xu, Jun Cheng, Avanti Shrikumar, Abhimanyu Banerjee, Daniel S Kim, Thorsten Beier, Lara Urban, et al. The kipoi repository accelerates community exchange and reuse of predictive models for genomics. Nature Biotechnology, 37(6):592–600, 2019.

[64] Fidel Ramírez, Devon P Ryan, Björn Grüning, Vivek Bhardwaj, Fabian Kilpert, Andreas S Richter, Steffen Heyne, Friederike Dündar, and Thomas Manke. deeptools2: a next generation web server for deep-sequencing data analysis. Nucleic Acids Research, 44(W1):W160–W165, 2016.

[65] Neva C Durand, James T Robinson, Muhammad S Shamim, Ido Machol, Jill P Mesirov, Eric S Lander, and Erez Lieberman Aiden. Juicebox provides a visualization system for hi-c contact maps with unlimited zoom. Cell Systems, 3(1):99–101, 2016.

